# Platelets regulate glioblastoma growth and immunity via sex-dependent PAR4 - Estrogen receptor beta signaling

**DOI:** 10.1101/2025.08.06.668464

**Authors:** Anthony R. Sloan, George Bukenya, Gavin P. Tannish, Daniel J. Silver, Anu Aggarwal, Juyeun Lee, Daniel Rosoff, Tyler J. Alban, Ivan Juric, Shikha Parsai, Vargab Baruah, Liza Tack, Tanvi Navadgi, Natalie Reitz, Shawn T. Ho, Aarav Badani, Jessica Goldberg, Xueer Yuan, Aymerick Gaboriau, Samrutha Kamatala, Jesse Coker, Vrishabhadev Sathish Kumar, Saachi S. Jain, Alliefair Scalise, Bhairavi Rajasekar, Alex Vincenti, Erin E. Mulkearns-Hubert, Craig M. Horbinski, Andrew E. Sloan, Christopher G. Hubert, Jingqin Luo, Joshua B. Rubin, Evi X. Stavrou, Falk W. Lohoff, Wendy A. Goodman, Tyler E. Miller, Christine M. O’Connor, Marvin T. Nieman, Naseer Sangwan, Timothy A. Chan, Alok A. Khorana, Andrew Dhawan, Scott J. Cameron, Justin D. Lathia

**Author notes:** **Corresponding author:** Justin D. Lathia, 9500 Euclid Ave, NE3-202 Cleveland, OH 44195.

## Abstract

Sex differences in cancer outcome, including that of glioblastoma (GBM), are shaped by biological, hormonal, and immunological factors and impact disease progression, treatment responses, survival, and tumor microenvironment (TME) interactions. Platelets regulate the immune responses and tumor progression in many cancers, but it is not clear how they contribute to these sex-based differences by affecting the dynamics of the TME. Here, we show that GBM patients exhibit heightened platelet reactivity driven by PAR4 signaling. In murine GBM models, both pharmacological inhibition of PAR4 using BMS986120 and genetic deletion of PAR4 significantly prolong survival in females but not males. This survival advantage is estrogen dependent: it is preserved in chromosomal male-hormonal female mice within the four-core genotype model and is rescued in ovariectomized mice treated with estrogen. The survival benefit is TME specific and is mediated by platelet-driven enhancement of CD8⁺ T cell infiltration into the tumor. Inhibition of platelet PAR4 signaling increases calcium signaling through an estrogen-dependent interaction between PAR4 and estrogen receptor β (ERβ)—a receptor interaction not previously described. PAR4-activated platelets within the TME suppress CD8⁺ T cell function, and depletion of CD8⁺ T cells abolishes both the tumor-induced platelet reactivity and the survival benefit conferred by PAR4 inhibition. These findings establish platelet-mediated PAR4 signaling as a critical driver of tumor progression and identify sex-specific immune responses as key to therapeutic efficacy.

## Introduction

Sex differences underlie and influence risk, progression, and treatment outcomes across many cancers. For example, men are more likely to develop several cancer types, including glioblastoma (GBM), while women often exhibit better survival rates^1, 2^. Despite these appreciated epidemiological differences, treatment paradigms remain uniform, highlighting the opportunity for sex-specific therapies shaped by hormonal, genetic, and immune differences. Sex differences in cancer also involve the tumor microenvironment (TME) — a complex network of cells and anatomical structures, including blood vessels, that interact with tumors. Among components of the TME, platelets play a major role in maintaining vascular integrity and modulating immune responses, potentially offering key insights into the relationship between sex differences and cancer progression. While platelets are present in the TME and known to influence immune cells, their role in cancer, particularly in thrombotic events and sex differences via thrombin-PAR4 signaling, remains underexplored^3–6^. In a recent study, the antiplatelet drug aspirin, which inhibits cyclooxygenase-1 and cyclooxygenase-2, enhanced the protective mechanism of T cells and therefore inhibited metastasis in murine models of lung and liver malignancies, further supporting the role of platelets in mediating cancer progression^7^.

In GBM, the most common primary malignant brain tumor in adults, immune cells in the TME suppress anti-tumor responses. Myeloid derived supporesor cells (MDSCs) drive immune suppression in GBM, while lymphocytes, such as CD8+ T cells, are reduced and have decreased T cell function^8–10^. Moreover, GBM has a striking difference in incidence and prognosis, with men 1.6 times more likely to develop the disease and exhibiting shorter survival^2, 11^. Both intrinsic factors and immune cells such as microglia, MDSCs, and CD8+ T cells contribute to these differences^8, 10^, with myeloid-derived cells having enhanced proliferation in males and male CD8+ T cells showing a greater propensity for exhaustion^10^. Although platelet-derived factors are known to promote tumor growth, their underlying mechanisms and role in sex-based differences in GBM are unclear^4, 12^. Thrombotic events, such as deep vein thrombosis (DVT) and pulmonary embolism (PE) are common in cancer patients and a leading cause of morbidity and mortality, although cause of death is not attributed to intravascular thrombosis^13, 14^. GBM patients face a 15-24% risk of thrombotic events, and patients with thrombotic events have worse outcomes, suggesting that a pro-thrombotic state drives cancer progression^15–18^. Therefore, given the sex difference in the incidence and aggressiveness of GBM that is driven by an immunosuppressive microenvironment more pronounced in males, we sought to determine whether sex-dependent platelet effects contribute to the immunosuppressive TME in GBM.

## Results

### PAR4 signaling drives hyperactive platelets in GBM patients

The GBM TME shapes the trajectory of tumor growth and response to therapies. A variety of mechanisms underlie TME-tumor interactions^1^; however, the role of platelets remains underappreciated. We and others have previously reported that high platelet counts are temporally related to worsened clinical outcomes in patients with any malignancy, including GBM^4, 19, 20^. By carefully evaluating a platelet transcriptomic signature in the context of large-scale human datasets, we found that an enrichment of a platelet gene signature is associated with poor overall survival in GBM patients (**Figure 1A, Figure S1A, Table S1**) and negatively correlated with several anti-tumor immune cell populations including T cells, dendritic cells (DCs), and B cells (**Figure S1B, Table S2**). These data indicates that platelets are a negative prognostic and immunosuppressive factor in GBM patients.

**Figure 1.**
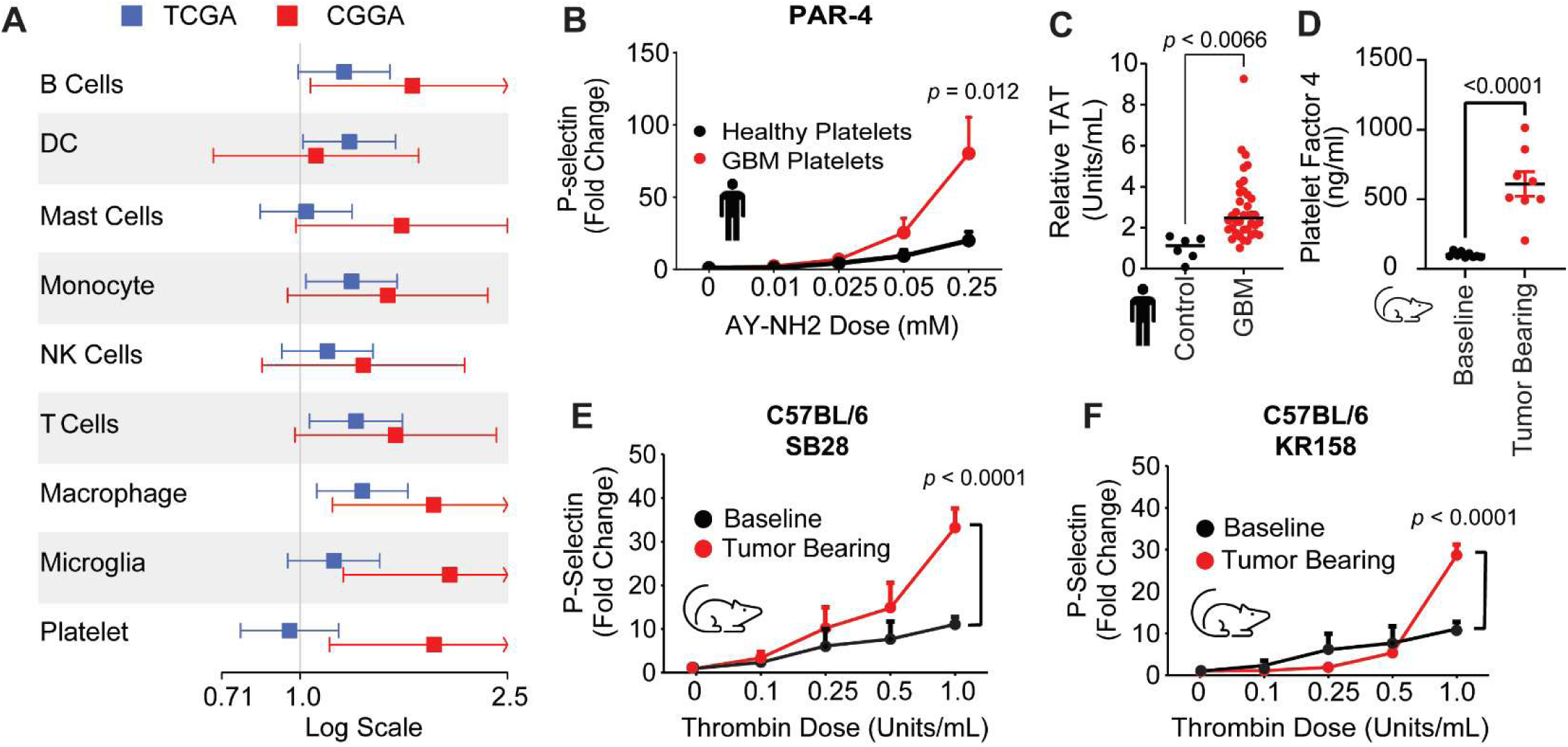
PAR4 signaling drives hyperactive platelets in GBM patients. **A**. WHO grade 4 IDH wildtype GBM samples from the publicly available data sets The Cancer Genome Atlas (TCGA) and Chinese Glioma Genome Atlas (CGGA_325). Univariate Cox proportional hazards models were constructed, and deconvoluted immune cell proportions and platelet signature score were z-score normalized and centered prior to hazard ratio computation. Hazard ratios depicted describe overall survival with 95% confidence intervals as a function of immune cell proportion or platelet signature score. **B**. Washed platelets isolated from patients with a GBM diagnosis and matched control subjects were treated with AY-NH2 to stimulate PAR4, and α-granule secretion was measured using an antibody specific for P-selectin via flow cytometry. Data are represented as means ± SEM from *n*=18 GBM patients and *n*=18 control subjects. Two-way ANOVA was performed. **C**. Thrombin-antithrombin (TAT) expression in plasma from GBM and healthy patient platelets as measured by TAT ELISA. An unpaired student’s *t*-test was performed. **D**. Serum was isolated from mice before intracranial tumor implantation (baseline) and following intracranial injection of the murine GBM SB28. Following tumor formation, platelet factor 4 (PF4 was measured via ELISA. An unpaired student’s *t*-test was performed. **E-F**. Washed platelets were isolated from mice before intracranial tumor implantation (baseline) and following intracranial injection of the murine GBM SB28 (**E**) and KR158 (**F**). Following tumor formation, washed platelets were isolated, and α-granule secretion was measured using an antibody specific for P-selectin via flow cytometry. Data are represented as means ± SEM from n=4 independent experiments. Two-way ANOVA was performed.

Next, we asked whether GBM patients have altered platelet physiology when compared to age- and sex-matched control subjects. We measured platelet reactivity from peripheral blood using assays that measure degranulation of platelet contents in GBM patients and control subjects (**Table S3, Table S4**). Platelets from both male and female GBM patients had increased agonist-induced reactivity, measured my P-Selectin, due to protease-activated receptor 4 (PAR4) signaling (**Figure 1B, Figure S2A-C**). However, agonist-induced platelet reactivity via protease-activated receptor 1 (PAR1), thromboxane receptor (TXA2), and purinergic receptor P2Y12 was similar or not as drastically different between GBM and control subjects (**Figure S2D-F**). Thrombin is a potent human platelet agonist that acts via the PAR1 and PAR4 receptors and is quantified by measuring plasma levels of thrombin-antithrombin (TAT) complexes. We found that GBM patients had significantly higher TAT levels relative to healthy age-matched control subjects (**Figure 1C**). Consistently, *in viv*o studies utilizing murine GBM models demonstrated that tumor-bearing mice had increased circulating platelet factor 4 expression (**Figure 1D**) and increased platelet reactivity compared to their baseline pre-tumor state (**Figure 1E, F**), with no difference in spontaneous or thrombin induced platelet reactivity between male and female mice (**Figure S2G-J**). Likewise, across patients with GBM, Using Mendelian randomization analyses, we found PAR4 expression is predictive of GBM diagnosis, whereas other platelet receptor expression was not predictive (**Figure S3A-C**). This observation is consistent with previous reports indicating that agonist-induced platelet activation and spontaneous platelet activation were both increased in patients with late-stage metastatic cancer^21^ and further suggest that tumor-induced platelet reactivity is mediated by PAR4.

### Inhibiting platelets via the thrombin-PAR4 signaling axis prolongs survival in a sex-dependent manner

To investigate the role of platelets in GBM tumor progression, we intracranially transplanted C57BL/6 mice with murine GBM cells followed by subsequent depletion of platelets through intravenous administration of anti-glycoprotein 1b (GP1b) antibody and survival analysis. These proof-of-concept studies demonstrated that platelet depletion prolongs survival in female tumor-bearing mice (**Figure 2A, Figure S4A**). Depletion of platelets had no effect in male mice in both tumor models (**Figure 2B, Figure S4B**), despite similar levels of platelet depletion between male and female mice (**Figure 2C, Figure S4C**). These results, in combination with the positive correlation between platelet expression and intratumoral immunosuppressive immune cell populations (**Figure 1A**), suggest that platelets drive tumor growth in a sex-biased manner, prompting us to question whether inhibiting the enhanced PAR4-mediated platelet reactivity may also show a sex difference in survival benefit.

**Figure 2.**
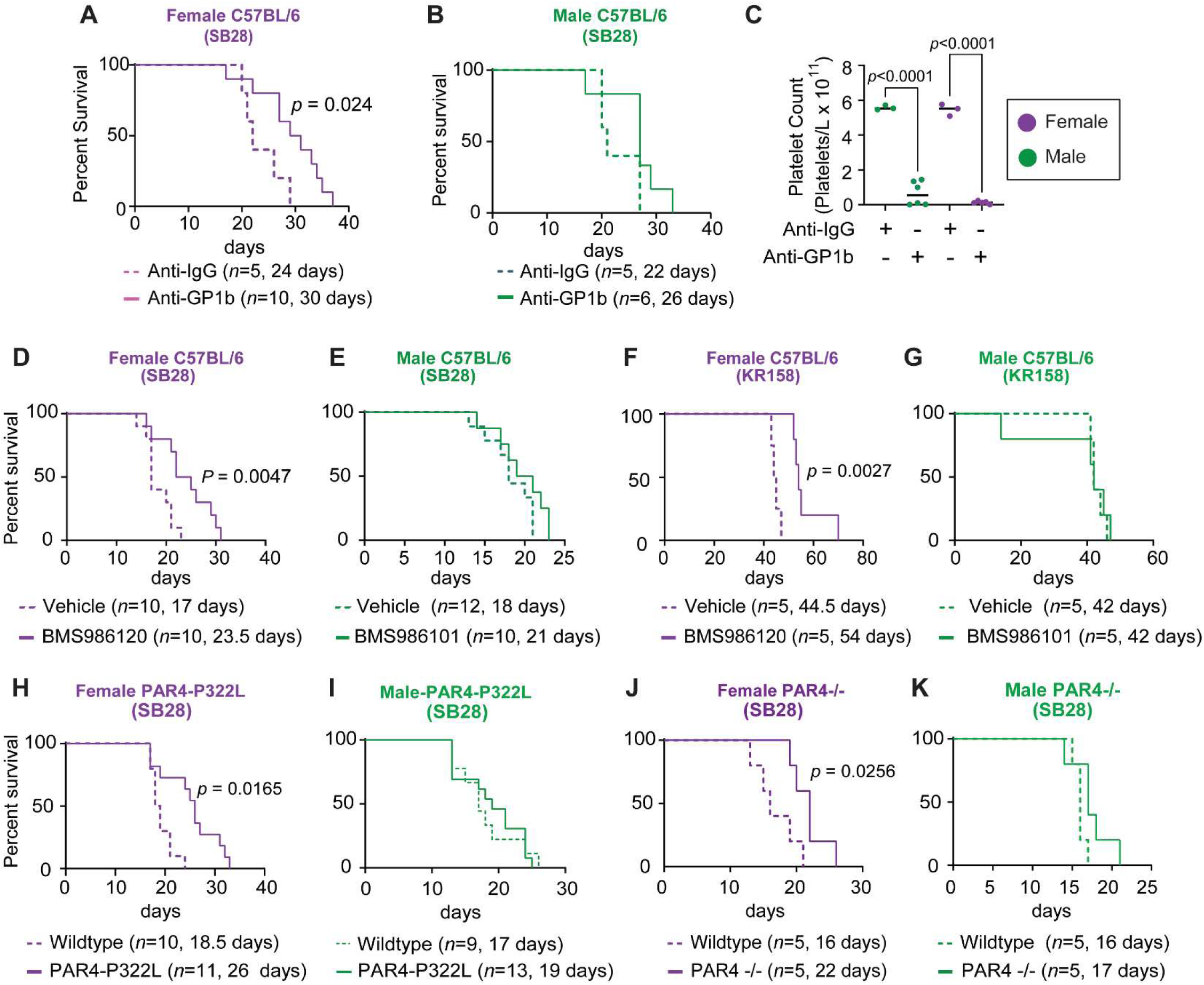
Inhibiting platelets via the thrombin-PAR4 signaling axis prolongs survival in a sex-dependent manner. **A**, **B**. Kaplan-Meier survival analysis was performed after intracranial tumor implantation of mouse GBM cell model SB28 in C57BL/6 female (**A**) and male (**B**) mice treated with anti-GP1bα depleting antibody. Statistical significance for survival analysis was determined by log-rank test. **C**. Complete blood count (CBC) platelet reading 1 day post anti-GP1bα treatment to validate platelet depletion in the tumor-bearing mice used in panels A and B. Data are shown with each point representing an individual animal. One-way ANOVA analysis with Tukey’s multiple comparison test was performed. **D-G**. Kaplan-Meier survival analysis was performed after intracranial transplantation of the mouse GBM cell models SB28 (**D, E**) and KR158 (**F, G**) in immunocompetent C57BL/6 mice followed by pharmacologically inhibiting the thrombin-PAR4 signaling axis by administering BMS986120 (2 mg/kg), a selective PAR4 inhibitor. Statistical significance for survival analysis was determined by log-rank test. **H**, **I.** Kaplan-Meier survival analysis comparing female (**H**) and male (**I**) C57BL/6 wildtype and C57BL/6 - PAR-P322L mice following intracranial transplantation of the mouse GBM cell model SB28. Statistical significance for survival analysis was determined by log-rank test. **J**, **K.** Kaplan-Meier survival analysis comparing female (**J**) and male (**K**) C57BL/6 wildtype and C57BL/6-PAR4-/- mice following intracranial transplantation of the mouse GBM cell model SB28. Statistical significance for survival analysis was determined by log-rank test.

We transplanted C57BL/6 mice with murine GBM cells and pharmacologically targeted the major platelet PAR receptor in mice (PAR4) with daily oral gavage treatment with the PAR4 antagonist BMS986120. Inhibiting PAR4 in tumor-bearing mice prolonged survival in a sex-dependent manner, with marked survival in females and no impact in males in all three murine GBM models tested (**Figure 2D-G**, **Figure S5A-B**). We further confirmed this sex-specific survival advantage using dabigatran, a direct inhibitor of thrombin, the PAR4 ligand (**Figure S5C-D**). To determine whether these effects were due to higher PAR4 expression in females, we examined intratumoral PAR4 (F2RL3) expression in GBM patient samples (TCGA) and found no difference in expression between tumors from males and those from females (**Figure S5E**). Subsequently, we measured surface and total PAR4 expression on mouse platelets and also found no difference in expression between males and females (**Figure S5F-G**). These studies suggest that inhibiting platelets via PAR4 results in prolonged survival in a sex-dependent manner.

To ascertain whether the effects observed were specific to PAR4 and not an off-target effect of BMS986120, we transplanted tumor cells into PAR4-P322L, which have a 70% reduction in PAR4 function^22^, and PAR4-/-transgenic animals. Consistently, we observed that female PAR4-P322L mice had a longer survival when compared to wildtype animals, whereas we observed no difference in male PAR4-P322L mice (**Figure 2H-I**). This was further confirmed in the PAR4-/- model, where female PAR4-/- mice had a longer survival compared to wildtype female animals, with no difference in male PAR4-/- mice (**Figure 2J-K**).

### Inhibiting the thrombin-PAR4 signaling axis prolongs survival in a female estrogen dependent manner

To investigate the impact of sex chromosomes, sex hormones, and their interaction on the female survival advantage when the thrombin-PAR4 signaling axis is targeted, we utilized the four-core genotype mouse model, which separates the effects of sex chromosomes from gonadal hormones. Using this model, we observed that chromosomal male (XY)-hormonal female mice exhibit a BMS986120-mediated survival advantage similar to that of wildtype females (**Figure 3A-B**), whereas chromosomal female (XX)-hormonal male mice do not (**Figure 3C-D**). These results suggest that the survival advantage seen upon BMS986120 administration is dependent on female sex hormones. To further validate this hormone dependency, we performed ovariectomies on female mice. One month later, we intracranially implanted GBM cells and administered BMS986120. Estrogen depletion rescued any prior survival benefit observed with BMS986120 administration compared to sham-operation control mice (**Figure 3E**). Finally, to determine whether this was estrogen specific, we utilized estrogen receptor-β knockout (ERβ⁻/⁻) mice to determine whether the female-specific survival benefit persisted in the absence of ERβ. Strikingly, genetic deletion of ERβ completely abolished the enhanced survival response to BMS986120 in female tumor-bearing mice (**Figure 3F**). These findings indicate that the female survival advantage is strictly ERβ-dependent and strongly support a model in which estrogen receptor signaling is required for the sex-specific effect of PAR4 inhibition.

**Figure 3.**
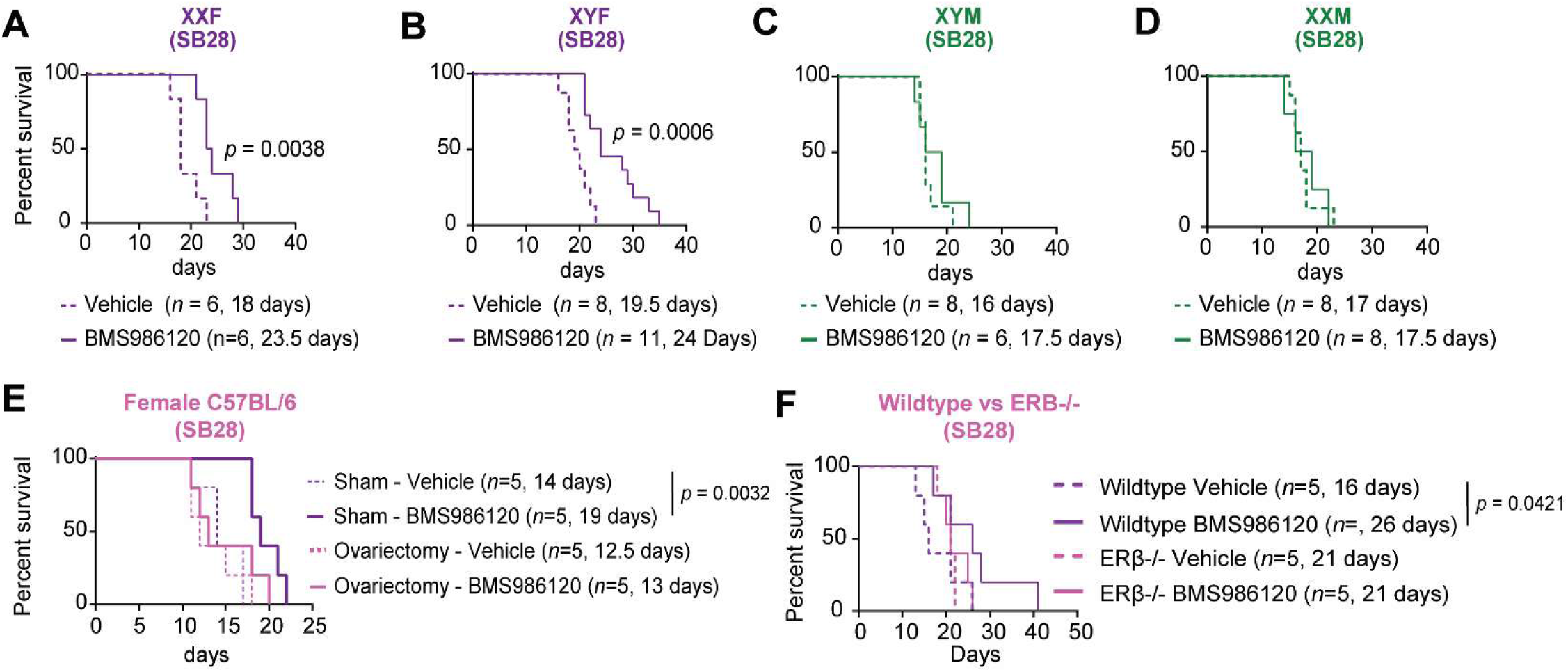
Inhibiting the thrombin-PAR4 signaling axis prolongs survival in a female estrogen-specific manner. **A-D.** Kaplan-Meier survival analysis was performed after intracranial transplantation of the mouse GBM cell model SB28 and treatment with BMS986120. The four core genotype model results in (**A**) wildtype female, (**B**) chromosomal male (XY)-hormonal female mice, (**C**) wildtype male, and (**D**) chromosomal female (XX)-hormonal male mice. Statistical significance for survival analysis was determined by log-rank test. **E**. Survival analysis of C57BL/6 female mice who underwent an ovariectomy or sham procedure and were intracranially transplanted with SB28 cells 30 days post ovariectomy and subsequently treated with BMS986120 (2 mg/kg). Statistical significance for survival analysis was determined by log-rank test. **F**. Kaplan-Meier survival analysis comparing female C57BL/6 wildtype and C57BL/6 – Erβ-/- mice following intracranial transplantation of the mouse GBM cell model SB28. Statistical significance for survival analysis was determined by log-rank test.

### Estrogen Receptor-β interacts with PAR4 and regulates PAR4 signaling

To determine whether the estrogen-dependent survival advantage is due to an estrogen response element on the *F2RL3* (PAR4-encoding gene) promoter, we treated pre-platelet megakaryocyte (MEG-01) cells with β-estradiol, dihydrotestosterone, or progesterone and explored *F2RL3* mRNA and PAR4 protein expression. We found that β-estradiol (**Figure S6A-B**), progesterone (**Figure S6C-D**), or dihydrotestosterone (**Figure S6E-F**) had no effect on *F2RL3* mRNA and PAR4 protein expression. This suggests that *F2RL3* mRNA and PAR4 protein expression are not altered by estrogen, progesterone, or androgen treatment, arguing against direct sex hormone regulation of the PAR4 promoter.

Given that the female-specific survival benefit of PAR4 inhibition was ERβ dependent, we next asked whether ERβ directly influences PAR4. To address this, we examined PAR4 expression and its interaction with ERβ. Our immunofluorescence imaging of MEG-01 cells revealed co-expression of PAR4 and Erβ (**Figure 4A**), suggesting these two receptors may physically interact. Indeed, lysates immunoprecipitated with ERβ and probed for PAR4 (**Figure 4B**) or the reciprocal (**Figure 4C**) revealed these proteins are in complex. This interaction was further confirmed by ERβ immunoprecipitation in human female platelets (**Figure 4D**) and human female brain microvascular endothelial cells (**Figure 4E**). These results suggest Erβ physically associates with PAR4 in platelets and endothelial cells, supporting a direct molecular interaction that may underlie the estrogen-dependent modulation of PAR4 signaling.

**Figure 4.**
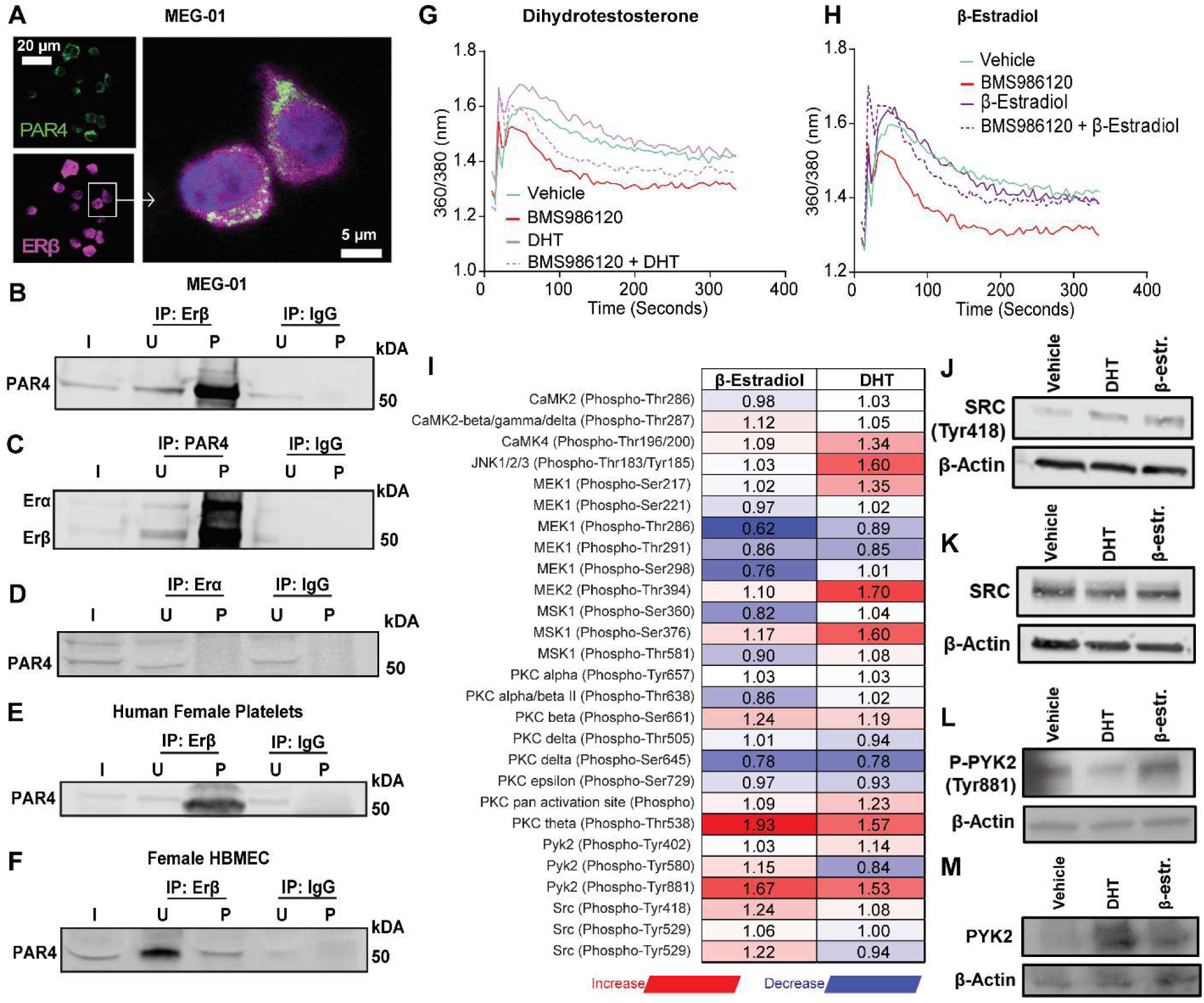
Estrogen receptor-β interacts with and regulates PAR4 signaling. **A.** Immunofluorescence (IF) staining of MEG-01 cells; IF staining of PAR4, Erβ. Merged image of DAPI, PAR4, and Erβ IF staining (Scale = 20 µm). **B.** Immunoprecipitation of Erβ and IgG followed by immunoblotting for PAR4 in MEG-01 cells. **C**. Immunoprecipitation of PAR4 and IgG followed by immunoblotting for Erβ in MEG-01 cells. **D**. Immunoprecipitation of Erα and IgG followed by immunoblotting for PAR4 in MEG-01 cells. **E.** Immunoprecipitation of Erβ and IgG followed by immunoblotting for PAR4 in human female platelets. F. Immunoprecipitation of PAR4 and IgG followed by immunoblotting for Erβ in human brain microvascular endothelial cells. **G**, **H**. Fura-2 calcium imaging of MEG-01 cells pretreated with BMS986120 and either (**G**) dihydrotestosterone or (**H**) β-estradiol followed by stimulation with thrombin. **I**. G protein-coupled receptor (GPCR) phospho-array of MEG-01 cells treated with β-estradiol or testosterone for 4 consecutive days. GPCR phospho-array was conducted and analyzed in collaboration with Full Moon Bio. Data represented as fold change relative to MEG-01 cells treated with vehicle. **J-M**. Validation of the GPCR phospho-array. MEG-01 cells were treated with β-estradiol or testosterone for 4 consecutive days, and lysates were immunoblotted for phospho-SRC (Tyr418) (**J**), SRC (**K**), phospho-PYK2 (Tyr881) (**L**), and PYK2 (**M**). Actin was used as a loading control.

Next, we evaluated PAR4 signaling and its inhibition by BMS986120 in the presence of β-estradiol to determine whether estrogen induces estrogen receptor-dependent regulation of PAR4. Knowing that calcium signaling is downstream of PAR4, we evaluated calcium. Using Fura-2 calcium imaging, we confirmed that PAR4 inhibition with BMS986120 suppresses Ca²⁺ signaling in MEG-01 cells, even when exposed to dihydrotestosterone (**Figure 4F**) or progesterone (**Figure S6G**). However, BMS986120 inhibition of PAR4 signaling was rescued when β-estradiol is present (**Figure 4G**). Additionally, utilizing a G-protein couple receptor (GPCR) phospho-array, we found that estrogen uniquely promotes coordinated activation of calcium-enriched signaling pathways in MEG-01 cells, a response not observed with testosterone. Estrogen engaged ERβ-dependent signaling marked by increased phosphorylation of SRC and PYK2, in addition to modulation of MAPK and PI3K signaling components (**Figure 4H-L**), recapitulating the established ERβ–SRC–PYK2–PI3K platelet signaling axis and overlapping with PAR4-dependent calcium regulation. Estrogen exposure induced coherent activation of calcium-responsive kinases, including CAMK2β/γ/δ and CAMK4, preservation of integrin-associated proteins, and selective activation of PKC isoforms (**Figure 4H-L**), consistent with sustained intracellular Ca²⁺ flux. These upstream events converged on Ca²⁺-linked MAPK intermediates, reflected by increased RSK and MSK phosphorylation, indicating engagement of the canonical CAMK-MAPK cascade. Together, these findings indicate that estrogen uniquely drives a calcium-dominant GPCR signaling program in megakaryocytes.

### Inhibiting thrombin-PAR4 signaling enhances CD8+ T cells in a sex-dependent manner

Next, we sought to determine the mechanism by which inhibiting the thrombin-PAR4 signaling axis prolongs survival in a sex-dependent manner. To determine whether the effect of BMS986120 was tumor cell intrinsic, we assessed proliferation in multiple murine GBM cell models treated with BMS986120 and observed no major differences in proliferation *in vitro* (**Figure S7A-B**), suggesting no significant impact on tumor cell fitness with BMS986120 treatment. These results led us to test whether the survival benefit seen in female tumor-bearing mice requires a fully functioning immune system. We intracranially transplanted NOD *scid* gamma (NSG) mice with murine GBM cells and subsequently inhibited PAR4 with BMS986120 administration. Pharmacologically inhibiting PAR4 with BMS986120 in immunodeficient mice did not show the survival benefit observed in immune-competent models (**Figure 5A, Figure S7C-D**). These findings reveal that the survival advantage observed in females when inhibiting the thrombin-PAR4 signaling axis depends on a functional immune system.

**Figure 5.**
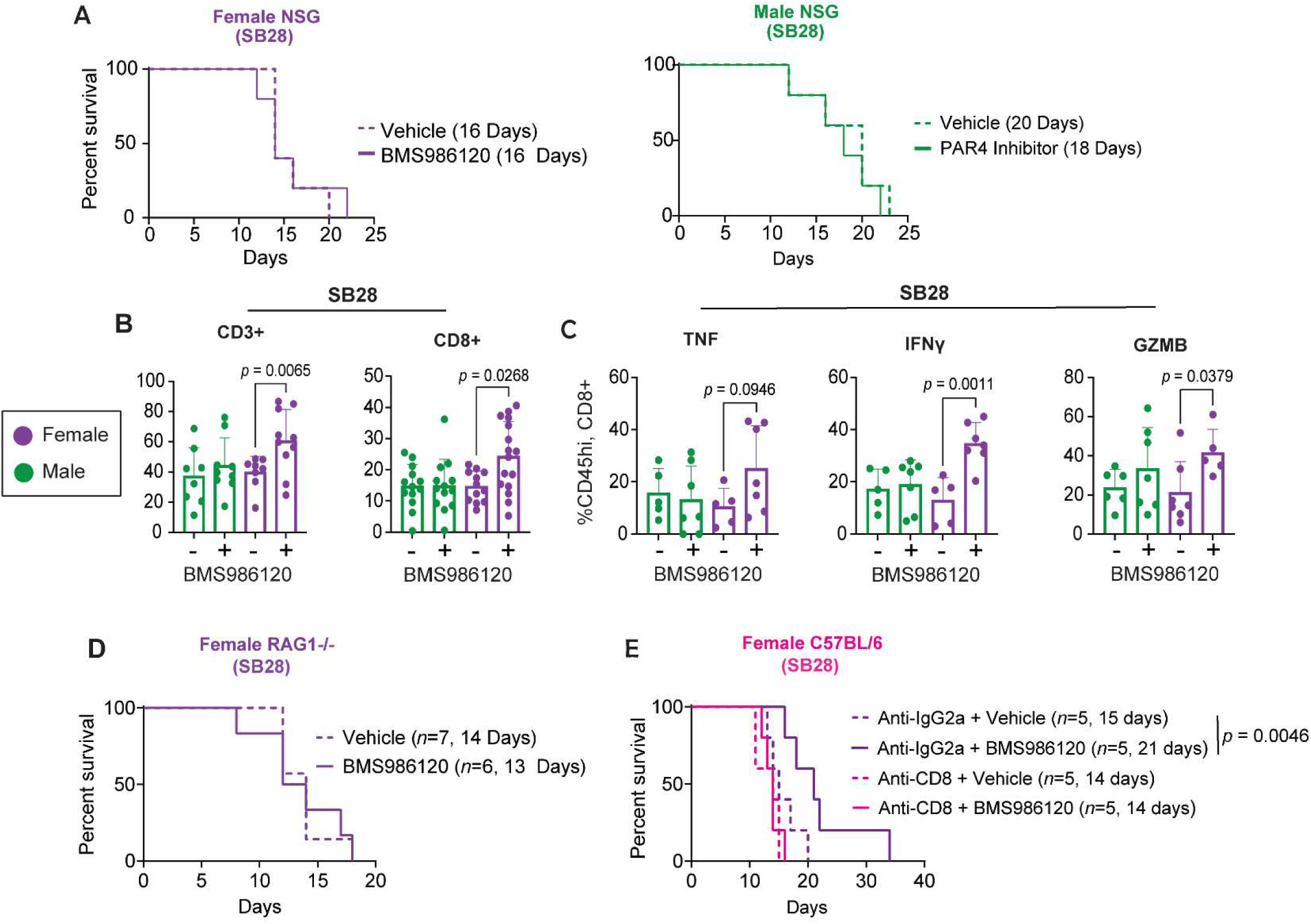
Inhibiting thrombin-PAR4 signaling enhances CD8+ T cells in a sex-dependent manner. **A.** Kaplan-Meier survival analysis was performed after intracranial injection of the mouse GBM cell model SB28 in immunodeficient NSG mice followed by pharmacologically inhibiting the thrombin-PAR4 signaling axis by administering BMS986120 (2 mg/kg), a selective PAR4 inhibitor. Statistical significance for survival analysis was determined by log-rank test. **B**. Frequency of tumor-infiltrating T-cell subsets 14 days post intracranial implantation of SB28 in male and female C57BL/6 mice treated with and without BMS986120. Data are combined from two independent experiments. One-way ANOVA analysis with Tukey’s multiple comparison test was performed. **C**. Intracellular cytokine expression in tumor-infiltrating CD8+ T cells. Data are combined from two independent experiments. One-way ANOVA analysis with Tukey’s multiple comparison test was performed. **D**. Kaplan-Meier survival curve depicting survival of SB28-bearing *Rag1-/-* female mice treated with BMS986120. Statistical significance for survival analysis was determined by log-rank test. **E**. Kaplan-Meier survival curve depicting survival of SB28 tumor-bearing females treated with anti-CD8-depleting antibody with BMS986120 administration. Statistical significance for survival analysis was determined by log-rank test.).

Platelets are known to influence immune cell function within the TME and in the systemic circulation^23–27^. While heterologous cell interactions are known to impact GBM, the question of whether platelets have sex-dependent effects is unexplored, and no studies have examined how PAR4 signaling regulates the TME across malignancies. To determine how the thrombin-PAR4 signaling axis contributes to the TME, we conducted immune profiling on tumor-infiltrating immune cells. We found that BMS986120-treated female mice had an increased percentage of CD45hi/CD3+ cells, specifically CD45hi/CD8+ tumor-infiltrating T cells 14 days post tumor implantation (**Figure 5B, Figure S7E, Table S5**), with no changes in other T cell populations or exhaustion markers (**Figure S8**). This BMS986120-mediated increase in CD8+ T cells was also seen at 10 days post BMS986120 treatment (**Figure S9, Table S5**), but not at day 7 (**Figure S10, Table S5**). Consistently, BMS986120-treated female CD8+ T cells had an increase in cytokine expression upon *ex vivo* stimulation at day 14 (**Figure 5C Table S6**) but not day 10 or day 7 (**Figure S9C-E**, **Figure S10C-E, Table S6**). There were no differences in other TME immune cell subsets (**Figure S11A-E, Figure S12A-H, Figure S13, Figure S14, Table S7**).

To further validate that the female survival benefit was CD8+ T cell dependent, we intracranially transplanted GBM cells into *Rag1-/-* mice, which lack T and B cells, and treated with BMS986120. Removing T cells reversed the survival benefit observed when mice were treated with BMS986120 (**Figure 5D**); however, when regulatory T cells (Tregs) were removed by treating tumor-bearing *FoxP3-DTR* mice with BMS986120, female mice still exhibited a survival advantage (**Figure S15**). To further isolate the effect specifically to CD8+ T cells, we intracranially implanted murine GBM cells into C57BL/6 mice and administered anti-CD8 depletion antibody in combination with BMS986120. Depletion of CD8+ T cells in the presence of BMS986120 completely rescued the female survival benefit observed when the thrombin-PAR 4 signaling axis was targeted (**Figure 5E, Figure S15B**). Lastly, using Akoya multiplex immunofluorescence imaging, we demonstrated that BMS986120 results in a decrease in intratumoral platelet expression (**Figure 6A-H, Figure S16A-D**) and an increase in CD8+ T cell expression in females, with no change in intratumor platelet or CD8+ T cell expression observed in male BMS986120 treated mice (**Figure 6I-N**). Distance proximity analysis between platelets and CD8+ T cells demonstrate trends where BMS986120 decreases the number of platelets in proximity to CD8+ T cells in both males and females (**Figure S16E**). Taken together, our results suggest that the survival benefit of inhibiting thrombin-PAR4 signaling in GBM is sex dependent, driven by sex hormones, and mediated through a decrease in platelet interactions with CD8+ T cells in the TME, highlighting a unique platelet-immune mechanism underlying therapeutic outcome.

**Figure 6.**
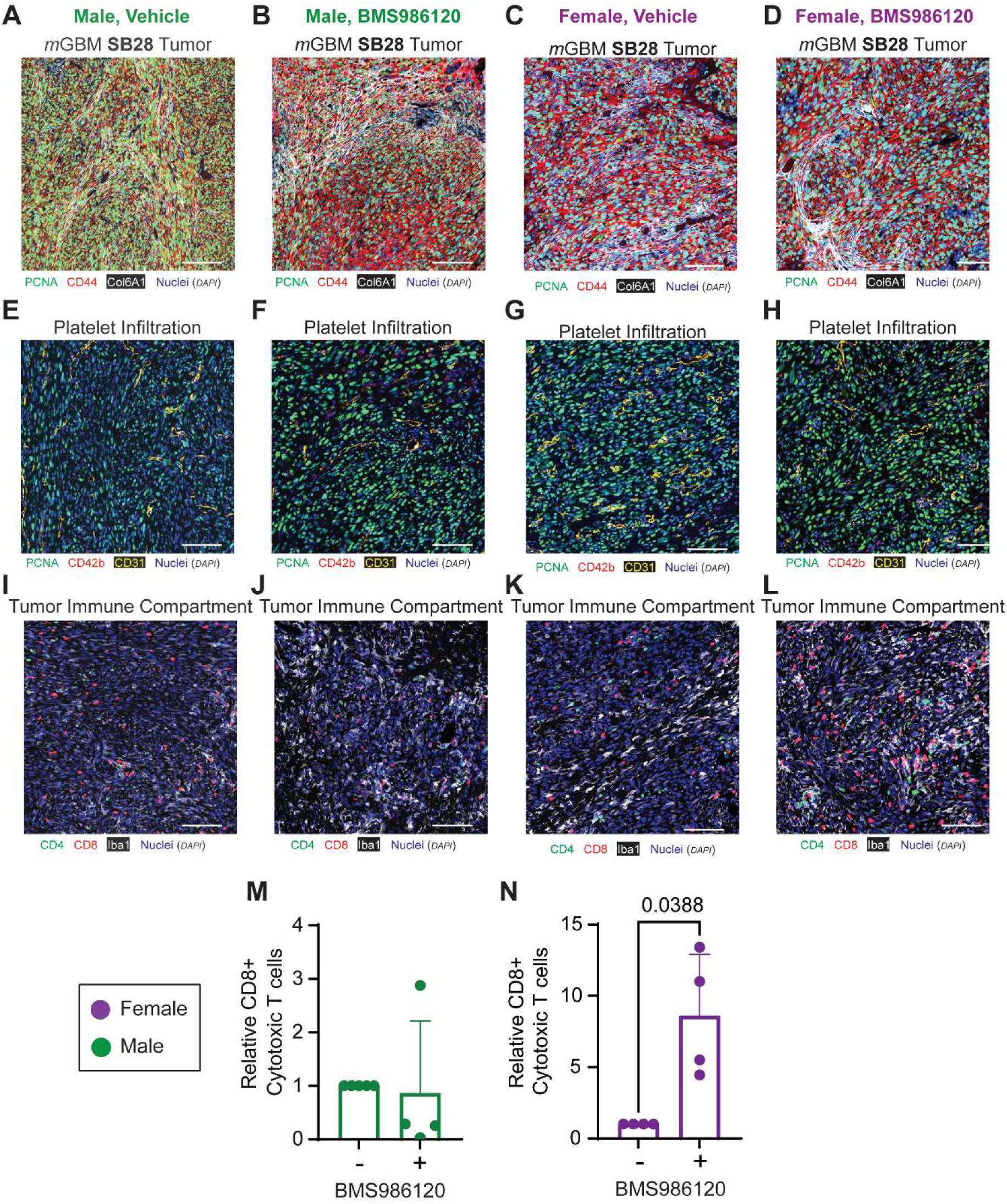
Inhibiting PAR4 increases intratumoral CD8+ T cells and decreases intratumoral platelets in tumor bearing mice. **A-D**. Akoya multispectral immunofluorescence imaging analysis of Merged image of DAPI, PCNA, CD44, and Col6A1 in (**A**) male vehicle treated, (**B**) male BMS986120 treated, (**C**) female vehicle treated, and (**D**) female BMS986120 treated. **E-H**. Akoya multispectral immunofluorescence imaging analysis of Merged image of DAPI, CD42b (platelet), CD31, and PCNA in (**E**) male vehicle treated, (**F**) male BMS986120 treated, (**G**) female vehicle treated, and (**H**) female BMS986120 treated. **I-L.** Akoya multispectral immunofluorescence imaging analysis of Merged image of DAPI, CD4, CD8, and Iba1 in (**I**) male vehicle treated, (**J**) male BMS986120 treated, (**K**) female vehicle treated, and (**L**) female BMS986120 treated. **M-N**. Akoya multispectral immunofluorescence quantification of CD8+ cytotoxic T cells from SB28 tumor bearing (**M**) male and (**N**) female vehicle and BMS986120 treated mice. An unpaired student’s *t*-test was performed.

### Inhibiting PAR4 enhances platelet secretion and Ca^2+^ signaling in an estrogen-dependent manner

PAR receptors, including PAR4, are thought to be expressed on platelets, endothelial cells and T cells^28^, implicating two potential mechanisms that could occur when PAR4 is targeted: a direct enhancement of CD8+ T cells in females or the inhibition of PAR4 signaling on platelets, which then indirectly enhances CD8+ T cell activity in females. To this end, we explored PAR expression in human and mouse T cells by utilizing publicly available datasets. We found that PAR4 was not expressed on T cells from mice (**Figure S17A-D)** or human (**Figure S18A-D)**. However, mice (**Figure S17A-D)** and human (**Figure S18A-D**) T cells did express high levels of PAR1. These results suggest that the increased tumor-infiltrating CD8+ T cells observed when inhibiting PAR4 in females is platelet dependent and that platelet signaling likely regulates tumor-infiltrating CD8+ T cells in a sex-dependent manner via PAR4.

We next determined the impact of pharmacologically inhibiting platelet PAR4 signaling on platelet function and signaling in tumor-bearing mice. We initially isolated platelets from male and female control and tumor-bearing mice and treated them with BMS986120 *ex vivo*. Using flow cytometry, we found no differences in p-selectin and activated GPllb/llla (JON/A) between tumor-bearing male and female platelets *ex vivo* (**Figure S19A, B**). These findings indicate that acute, *ex vivo* pharmacologic inhibition of PAR4 does not intrinsically alter platelet activation or α-granule secretion in a sex-dependent manner, suggesting that the sex-specific effects of PAR4 inhibiting observed in vivo likely require the tumor microenvironment or sustained systemic signaling rather than direct platelet-intrinsic differences alone. Next, we intracranially transplanted GBM cells and treated mice with BMS986120 for 14 days. Aggregation assays demonstrated that pharmacologically inhibiting PAR4 for 14 days had no effect on aggregation/thrombosis in tumor-bearing mice (**Figure 7A**). However, flow cytometry for p-selectin on platelets showed that female tumor-bearing mice had inhibited α-granule secretion following a 14-day treatment with BMS986120 (**Figure 7B, Figure S19C**), whereas male mice treated with BMS986120 had no change in alpha granule secretion (**Figure 7C, Figure S19D**).

**Figure 7.**
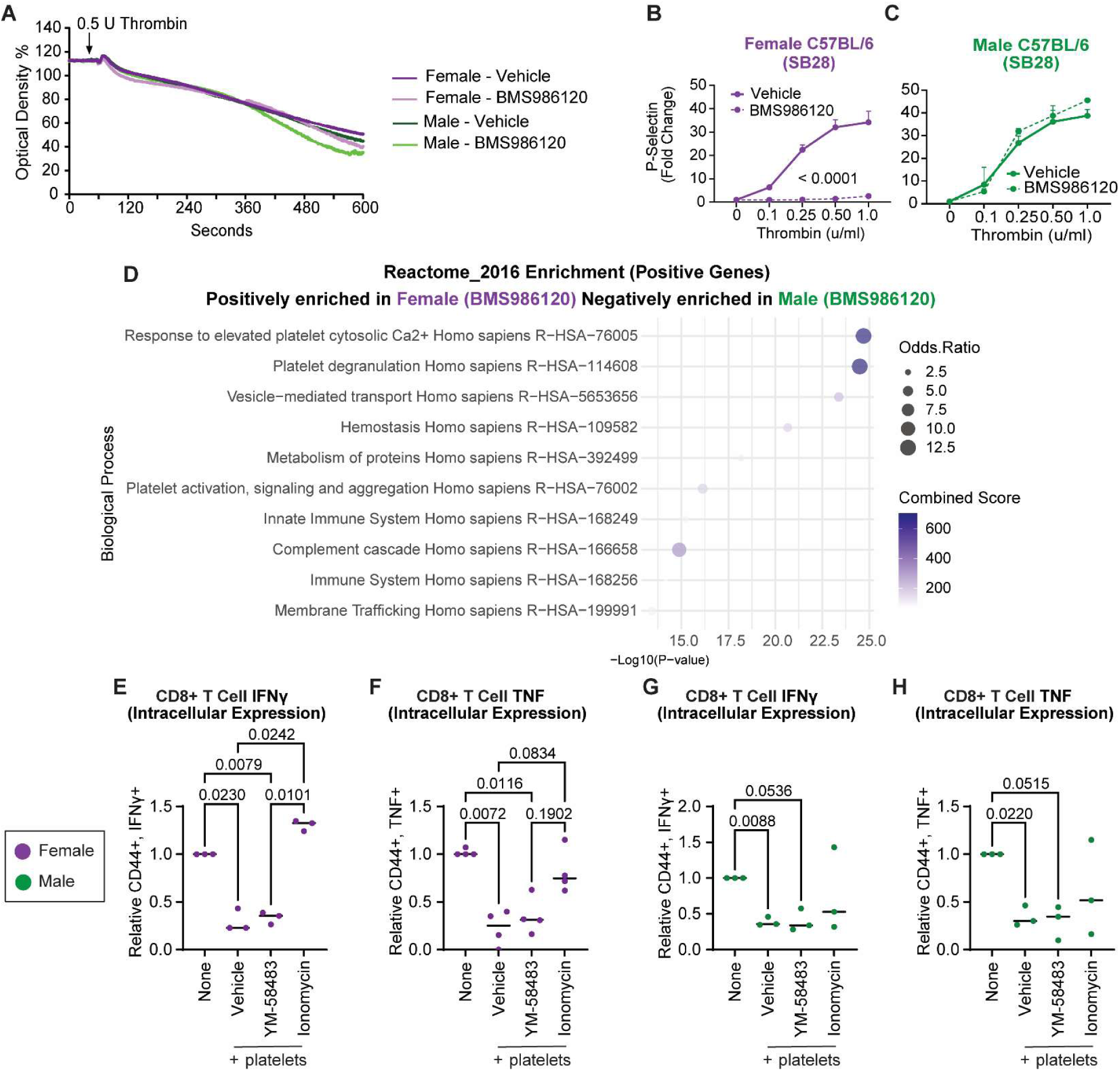
Inhibiting PAR4 enhances Ca^2+^ signaling, platelet secretion and exocytosis in a sex-dependent manner. **A.** Aggregometry analysis of washed platelets isolated from mice treated with BMS986120 following intracranial implantation of SB28 and stimulated with 0.5 U thrombin. Statistical significance was determined by a two-way ANOVA. **B, C.** Washed platelets isolated from mice treated with BMS986120 following intracranial implantation of SB28. Washed platelets were isolated, and α-granule secretion was measured using an antibody specific for P-selectin. Data are represented as means ± SEM. Statistical significance was determined by two-way ANOVA. **D**. Proteomic GO enrichment analysis of washed platelets isolated from mice treated with BMS986120 following intracranial implantation of SB28. Pathways are positively enriched in female BMS986120 platelets and negatively enriched in male BMS986120 platelets. **E-H**. Intracellular cytokine expression of CD8+ T cells isolated from (**E**, **F**) female and (**G**, **H**) male C57BL/6 mouse spleens and co-cultured with sex-matched washed platelets isolated from B6 wildtype mice; platelets were pre-treated with vehicle, YM-58483, or ionomycin before addition to platelets, fresh platelets were added for 3 consecutive days, and intracellular IFNγ (**E, G**) and TNF (**F**, **H**) expression was measured by flow cytometry. One-way ANOVA analysis with Tukey’s multiple comparison test was performed.

Proteomic GO enrichment analysis demonstrated that peripheral platelets from BMS986120-treated female mice had enriched platelet degranulation and exocytosis signaling pathways (**Figure 7D**). Additionally, the most highly enriched pathway in BMS986120-treated female mice was elevated platelet cytosolic Ca^2+^. Consistently, GO enrichment analysis on vehicle-treated male and female tumor-bearing mice showed differences in several transport and exocytosis pathways, with no difference in elevated platelet cytosolic Ca^2+^ between male and female untreated platelets (**Figure S19E**).

Calcium is essential for CD8⁺ T-cell activation, promoting their recruitment, cytokine production, and cytotoxic function through Ca²⁺-dependent signaling pathways^29^. To address whether the enhanced Ca^2+^ signaling in female platelets when inhibiting PAR4 with BMS986120 is responsible for increased CD8+ T cell function in the TME, we pre-treated platelets with the calcium release-activated calcium (CRAC) channel blocker YM-58483 or the calcium ionophore ionomycin. We found that pre-treating platelets with YM-58483, inhibited cytokine expression similar to that of platelets; however, pre-treating platelets with ionomycin enhanced cytokine expression in CD8+ T cells (**Figure 7E-H**), with a more pronounced effect in females (**Figure 7E-F**) than in males (**Figure 7G-H**). Together, these results suggest that PAR4 inhibition enhances tumor-infiltrating CD8⁺ T cells through a platelet-intrinsic, sex-dependent mechanism in which pharmacologic inhibition of PAR4 selectively alters calcium signaling and α-granule secretion in female platelets without affecting global aggregation.

### Bidirectional regulation of functions between platelets and CD8+ T cells in GBM

Our observations and previous work have demonstrated that high platelet counts associate with poor survival in GBM and multiple other malignancies^4, 30–33^. Platelets also interact with and educate the function of other cells, including immune cells that have been identified in the TME and within the circulation^23–27^. However, the role of platelets in regulating the immunosuppressive TME remains poorly understood. When CD8+ T cells were co-cultured with fresh platelets (provided daily), directly or using a transwell, T cell function as measured by intracellular cytokine expression (**Figure 8A-B, Figure S20A-B**) and cytokine secretion (**Figure 8C-D**), was inhibited. When PAR4 function was decreased to 60% by using platelets from PAR4-P322L mice, this observed effect was reversed in females but not male mice (**Figure 8A-B**). Additionally, inhibiting PAR4 with BMS986120 reversed the effect of platelets on cytokine secretion in females but not males (**Figure 8C-D**). In an *in vivo* setting, GBM tumor-bearing immunocompetent mice have constitutively higher platelet reactivity relative to their baseline. However, platelet hyperreactivity was not observed in tumor-bearing NSG mice, which lack T, B, and NK cells (**Figure S20C**). To further decipher whether tumor-induced platelet reactivity is T cell specific, GBM tumors were induced in *Rag1-/-* mice, which have no B or T cells. In this case, no increase in platelet reactivity was observed (**Figure 8E**). Consistently, we also tested platelet activity in tumor-bearing mice that were pretreated with an anti-CD8 antibody, and no enhanced platelet reactivity was found (**Figure 8F**). These results reveal that tumor-induced platelet reactivity in murine GBM models is dependent on the presence of immune cells, particularly CD8+ T cells, and that subsequent CD8+ T cell inhibition of platelets is dependent on PAR4 (**Figure 8G**).

**Figure 8.**
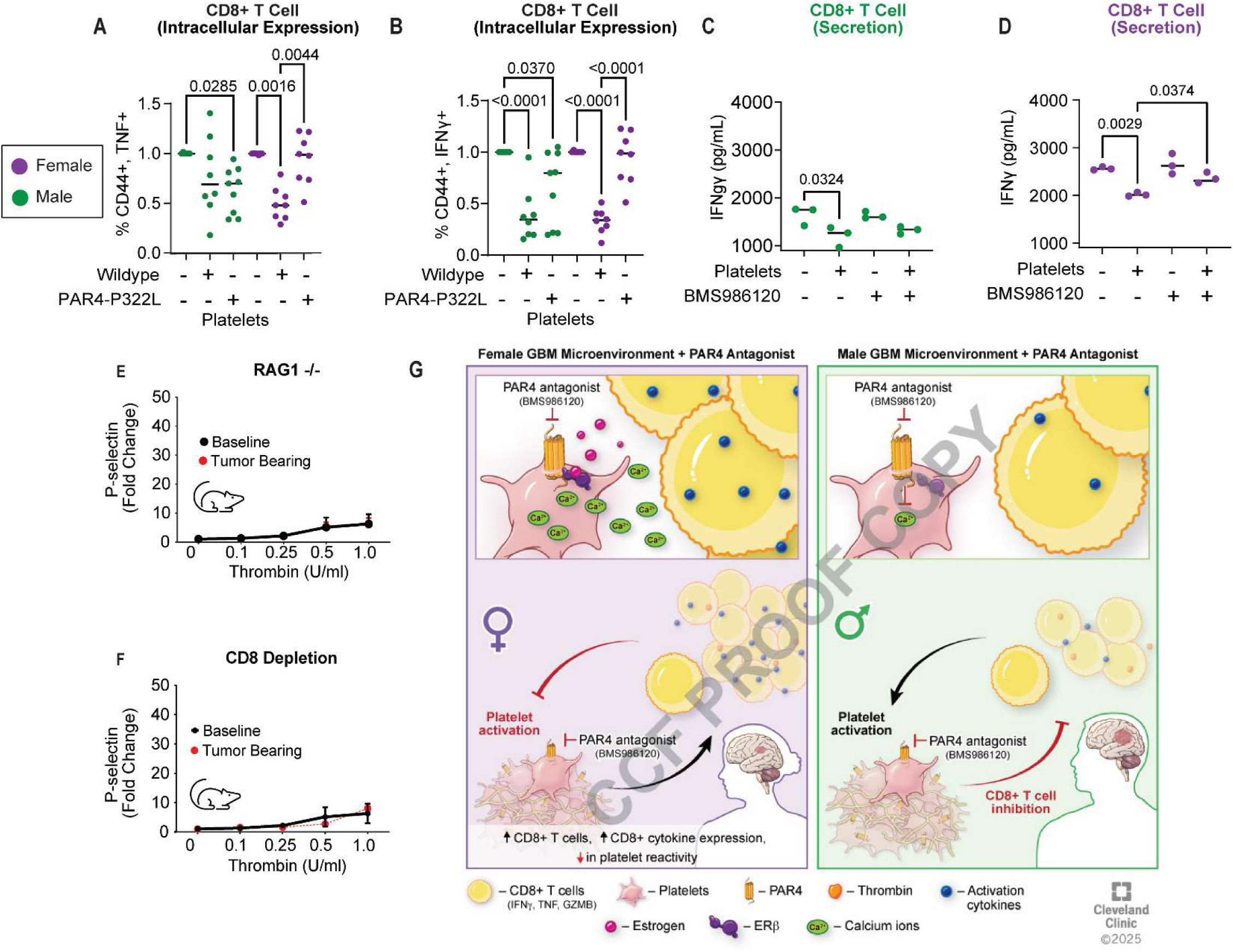
Bidirectional regulation between platelets and CD8+ T cells in GBM. **A, B**. Intracellular cytokine expression of CD8+ T cells isolated from C57BL/6 mouse spleen and co-cultured with sex-matched washed platelets isolated from C57BL/6 mice or PAR4-P322L mice; fresh platelets were added for 3 consecutive days, and intracellular TNF (**A**) and IFNγ (**B**) expression was measured by flow cytometry. One-way ANOVA analysis with Tukey’s multiple comparison test was performed. **C, D**. IFNγ cytokine secretion from male (**C**) and female (**D**) CD8+ T cells isolated from C57BL/6 mouse spleen and co-cultured with sex-matched washed platelets isolated from C57BL/6 mice; fresh platelets were added for 3 consecutive days, and supernatant IFNγ secretion was measured by ELISA. One-way ANOVA analysis with Tukey’s multiple comparison test was performed. **E, F**. Washed platelets isolated from *Rag1-/-* mice (**E**) and C57BL/6-CD8-depleted mice (**F**) before intracranial tumor implantation (baseline) and two weeks following intracranial injection of SB28 cells. Washed platelets were isolated, and α-granule secretion was measured using an antibody specific to P-selectin via flow cytometry. Data are represented as means ± SEM from n=4 independent experiments. **G**. Proposed model where platelet-mediated PAR4 signaling is as a critical driver of tumor progression and identifying sex-specific immune responses as key to therapeutic efficacy.

## Discussion

Our study identifies a novel connection between platelet signaling, hormonal regulation, and immune activity within the GBM TME. Here, we show that GBM patients exhibit heightened platelet reactivity driven specifically by PAR4 signaling. In murine GBM models, both pharmacological inhibition of PAR4 using BMS986120 and genetic deletion of PAR4 significantly prolonged survival in females but not males. This survival advantage is estrogen dependent: it is preserved in chromosomal male–hormonal female mice within the four-core genotype model and is rescued in ovariectomized mice treated with estrogen. The survival benefit is TME specific and is mediated by platelet-driven enhancement of CD8⁺ T cell infiltration into the tumor. Inhibition of platelet PAR4 signaling increases calcium signaling through an estrogen-dependent interaction between PAR4 and estrogen receptor β (ERβ), a receptor interaction not previously described.

Although BMS986120 reduced platelet–CD8⁺ T cell proximity in both males and females, only females demonstrated a corresponding increase in intratumoral CD8⁺ T cell accumulation. We speculate that this divergence reflects estrogen–estrogen receptor β signaling in female platelets, which may uniquely couple thrombin–PAR4 inhibition to enhanced CD8⁺ T cell recruitment and/or retention within the tumor microenvironment. PAR4-activated platelets within the TME suppress CD8⁺ T cell function, and depletion of CD8⁺ T cells abolishes both the tumor-induced platelet reactivity and the survival benefit conferred by PAR4 inhibition (**Figure 8G**). This is achieved through perturbations in immune function within the TME, leading to sex bias in immune cell subpopulations such as microglia, MDSCs, and T cells^8–10^. This study highlights cancer-associated thrombosis as a key driver of tumor progression, supporting precision medicine strategies that account for sex as a biological variable.

Cancer patients have a significantly higher risk of developing thrombotic events compared to the general population, with an estimated ninefold increase in venous thromboembolism risk^13, 14, 34, 35^. Recent studies highlight the role of platelet activation in the TME, where they influence immune responses and promote pro-thrombotic conditions^7^. One likely source of thrombin in the GBM TME is the tumor cells themselves: we previously showed that cancer stem cells (CSCs) can endogenously produce all coagulation cascade components and generate biologically active thrombin locally — independent of plasma — providing a direct source within the tumor. In addition, thrombin in the TME could arise from: (1) expression of procoagulant factors (e.g., tissue factor) by tumor or stromal cells that drive local coagulation ^16, 36, 37^; (2) disturbed or leaky tumor-associated vasculature, leading to plasma protein extravasation and local activation of coagulation; or (3) pro-coagulant extracellular vesicles or microparticles released by tumor, endothelial, or immune cells that catalyze thrombin generation. The broader phenomenon of tumor-driven coagulation activation via tissue factor and coagulation cascade components has been well described in solid cancers (and brain tumors).

Our work expands on this concept by demonstrating that tumor formation heightens platelet reactivity through an additional platelet receptor, PAR4, which directly contributes to immune suppression in a sex-dependent manner. This underscores how tumors hijack normal physiological processes to create a microenvironment that supports their progression and survival. Our findings emphasize the importance of inhibiting cancer-associated thrombosis and platelet activation and suggest that, with further study, selective anticoagulation strategies might complement existing therapies and potentially enhance antitumor responses. Additionally, although this study focuses on GBM, the mechanistic coupling among platelet PAR4 signaling, thrombotic pathways, and immune modulation suggests broader relevance across cancers with high thrombotic burden. In addition to triple-negative breast cancer, these pathways may be particularly relevant in estrogen receptor-positive malignancies, where estrogen signaling could further intersect with platelet activation and immune regulation. Together, these observations raise the possibility that sex- and hormone-dependent PAR4 signaling may influence antitumor immunity across multiple cancer types, including pancreatic, colon, and lung cancers, warranting future investigation.

Inhibiting PAR4 offers a therapeutic advantage over inhibiting PAR1 due to its distinct signaling kinetics, which reduce bleeding risks while still effectively modulating platelet activity^22, 38, 39^. Unlike PAR1, which responds rapidly to low thrombin concentrations, PAR4 requires higher thrombin levels for activation, leading to more highly controlled platelet responses and minimizing excessive bleeding complications. Additionally, PAR4 mediates the prolonged signaling necessary for stable thrombus formation, whereas PAR1 activation results in a rapid, transient signal. By selectively inhibiting PAR4, therapeutic strategies can limit thrombosis while preserving essential hemostatic functions, reducing the likelihood of severe bleeding events associated with PAR1 inhibition^22^. Previous work has demonstrated that inhibiting PAR4 enhances CD4+ T regulatory cells, contributing to immunosuppression^40^; however, our results show that PAR4 inhibition enhances CD8+ T cell function in a sex-dependent manner. This could be because T cells in brain tumors often exhibit exhaustion, immunosuppression, and altered metabolism, which weaken their ability to effectively combat the tumor^41^. Compared to other tumor types, T cells in brain tumors face heightened challenges due to the brain’s unique immune-privileged status and the suppressive tumor microenvironment, which includes factors like regulatory T cells and tumor-associated macrophages^42^. These elements contribute to reduced T cell infiltration and impaired antitumor responses. These findings highlight PAR4 as a more favorable target for platelet inhibition in cancer-associated thrombosis and other thrombotic disorders in females.

While our findings establish the role of PAR4-mediated platelet signaling in GBM progression and its sex-dependent therapeutic implications, several limitations remain. Translating results from murine models to human patients requires further clinical validation across diverse genetic and tumor backgrounds. Additionally, while estrogen contributes to the survival benefit of PAR4 inhibition, broader hormonal influences on platelet function and immune modulation warrant further exploration. An important consideration not addressed in the present study is the impact of cyclical estrogen fluctuations on PAR4 expression and therapeutic response. Although hormone cycle-dependent regulation of PAR4 could have implications for optimizing the timing of PAR4 antagonist administration, incorporating estrous cycle–specific analyses in an intracranial GBM model presents substantial technical and logistical challenges, including longitudinal hormone monitoring and repeated tissue sampling. Future studies specifically designed to integrate hormone-cycle staging with PAR4 expression and drug timing will be critical to determine whether estrogen-dependent dynamics can be leveraged to inform personalized dosing strategies for PAR4-targeted therapies. Additionally, the therapeutic potential of BMS986120 is promising, but the long-term efficacy, safety, and off-target effects must be assessed in preclinical and clinical models.

Based on our findings, future research is well positioned to investigate alternative platelet secretion mechanisms in shaping immune responses, with proteomic analyses identifying sex-specific signaling pathways that influence treatment efficacy. Examining how estrogen affects platelet-driven TME dynamics may yield novel hormone-modulating strategies to complement PAR4-targeted therapies. Expanding these insights into human studies and developing biomarkers for platelet activity could better predict patient outcomes. Our findings highlight the importance of addressing cancer-associated thrombosis and platelet activation as potential therapeutic targets and suggest that future studies should explore how anticoagulation strategies might interact with immune processes in GBM to improve patient outcomes.

## Materials and Methods

### Cell models

The syngeneic mouse GBM cell line SB28 was generously gifted by Dr. Hideho Okada (University of California San Francisco), and KR158 was generously gifted by Dr. Loic P. Deleyrolle (Mayo Clinic, previously University of Florida). The GL261 line was from the Developmental Therapeutics Program, NCI. MEG-01 cells were generously gifted by Dr. Scott J. Cameron (Cleveland Clinic Research). All cell lines were routinely tested for *Mycoplasma* and were preemptively treated with 1:100 MycoRemoval Agent (MP Biomedicals) upon thawing. All cell lines used in these studies were maintained in complete RPMI 1640 (Media Preparation Core, Cleveland Clinic Research) supplemented with 10% FBS (Thermo Fisher Scientific), 1% penicillin/streptomycin (Media Preparation Core, Cleveland Clinic Research). All cell lines were maintained at 37°C and 5% CO_2_ in a humidified incubator.

### Mice

All animal experiments were performed in accordance with Cleveland Clinic IACUC policies and guidelines and approved by the IACUC committee at Cleveland Clinic Research. C57BL/6 (RRID:IMSR_JAX:0006641), *Rag1^-/-^* (B6.129S7-*Rag1^tm1Mom^/J*; RRID:IMSR_JAX:002216), four-core genotye (FCG) (B6.Cg-Tg(Sry)2Ei *Sry^dl1Rlb^*/JtuJ), and NOD.Cg-*Prkdc^scid^ Il2rg^tm1Wjl^*/SzJ (NSG; RRID:IMSR_JAX:005557) mice were purchased from The Jackson Laboratory as required. PAR4-P322L and PAR4-/- mice were a generous gift from Dr. Marvin Nieman at Case Western Reserve University^22^. Estrogen receptor beta knockout (Erβ-/-) mice were a generous gift from Dr. Wendy Goodman at Case Western Reserve University.

### Intracranial tumor transplantation and treatments

For intracranial tumor implantation, 4- to 7-week-old mice were intracranially injected with 10,000 to 20,000 cells in 10 µL of RPMI-null media, 3 mm caudal to the coronal structure, 1.8 mm lateral to the sagittal suture at a 90° angle with the skull to a depth of 3.0 mm. Animals were monitored daily for neurological symptoms that include lethargy and hunched posture to qualify signs of tumor burden. In some experiments, male and female mice were treated with 2 mg/kg BMS986120 (MedChemExpress) or vehicle (8% DMSO) via oral gavage, daily, beginning 1-7 days post tumor implantation depending on the tumor latency of the model. In other experiments, to deplete CD8+ T cells, mice were intraperitoneally injected with an anti-CD8 antibody (BioXCell, Cat. #0061) or isotype control antibody (BioXCell, cat. #BE0090). Animals were administered 200 µg of antibody on day −1, followed by 100 µg every 5 days until humane endpoint was reached as previously described^10^. In other experiments, to deplete platelets, anti-GPlbα (EMFRET, Cat. R300) or anti-igG was administered intraperitoneally 1 day post tumor implantation to avoid hemorrhage at a concentration of 1 µg/g. Subsequent administration was performed at a concentration of 0.5 µg/g every 5 days until humane endpoint was reached.

### Ovariectomy

For oophorectomy procedures, 4- to 7-week-old female mice were deeply anesthetized, and a longitudinal incision was made 0.5 cm above where the upper border line of the two hind limbs connect with the back. The skin incision was pulled approximately 0.5 cm horizontally to the left or the right to visualize two soft, white, shiny fat masses next to the lower pole of the kidneys. The left and right ovaries of the mouse are found in the respective fat masses and were bi-laterally ovariectomized by cutting each side of the ovary and excising with sterile surgical scissors, which results in little to no bleeding. After the muscle layer was closed with sutures and the skin incision was sutured or stapled, the mice were placed in a heated chamber to recover. Mice in the sham group also underwent a similar procedure as the ovariectomized group, except the ovaries were left intact and only some adipose tissue near the ovaries was removed.

### Human platelet isolation and stimulation

Human platelet isolation was conducted as previously described^43, 44^. In brief, whole blood was collected from healthy donors into tubes containing 3.2% sodium citrate (BD Vacutainer) using a 21-gauge butterfly needle to ensure minimal activation during collection. Platelet-rich plasma (PRP) was obtained by centrifuging the collected blood at 200 x g for 15 minutes at room temperature, carefully avoiding the buffy coat. PRP was then subjected to a second centrifugation at 200 x g for 15 minutes to further purify the sample. The resulting supernatant was discarded, and the PRP was diluted 1:1 with Tyrodes buffer. Prostaglandin I2 (PGI2, 10 nM final concentration) was added to prevent spontaneous platelet activation during the isolation process. The PRP-Tyrode’s mixture was centrifuged at 1,400 x g for 5 minutes to pellet the platelets, and the supernatant was removed. Washed platelets were resuspended in Tyrode’s buffer and diluted 1:10 to achieve a working platelet suspension. Fifty million platelets per mL was used immediately for subsequent activation assays.

For platelet activation, aliquots of 100 µL of the platelet suspension were stimulated with increasing concentrations of agonists inhibiting the PAR1 receptor (TRAP6), the thromboxane receptor (U46619), the P2Y12 receptor (ADP), and PAR4 (AY-NH2). Agonist concentrations ranged from 0.01 µM to 20 µM for TRAP6, U46619, and ADP, and from 0.05 µg/mL to 1 µg/mL for CRP. After a 15-minute stimulation at room temperature, 1 µL of P-selectin antibody (anti-CD62P, clone AK-4, eBioscience) conjugated to PE was added to each sample and incubated in the dark for 30 minutes to prevent photobleaching.

Following antibody staining, 100 µL of 2% formalin was added to each sample to fix the platelets. Fixed samples were stored at 4°C for no longer than 16 hours prior to analysis. Flow cytometry was performed on a BD Accuri C6, and fluorescence was detected using the FL-2 channel (for PE-conjugated antibodies). Platelet activation was quantified based on the surface expression of CD62P, with data analyzed using FlowJo software.

### Mouse platelet isolation and stimulation

Whole blood was collected from C57BL/6 mice via retro-orbital bleeding using heparinized capillary tubes. Blood was drawn into tubes containing heparin/Tyrode’s solution (1000 U/mL heparin in Tyrode’s buffer, 1:20 dilution). The collected blood was gently mixed by inverting the tube and processed immediately. PRP was isolated by centrifuging the whole blood at 1000 RPM for 5 minutes at room temperature. Plasma was carefully collected, avoiding red blood cells, and subjected to a second centrifugation at 1000 RPM for 5 minutes to remove remaining blood cells. The PRP was then supplemented with 1 µM PGI2 to prevent platelet activation during isolation. The platelet pellet was obtained by centrifuging PRP at 2700 RPM for 5 minutes, after which the supernatant was discarded. Platelets were resuspended in Tyrode’s buffer and pooled to achieve a final concentration suitable for stimulation experiments. Platelets were aliquoted into 96-well plates at a concentration of 100 µL per well for subsequent activation assays.

For platelet activation, 1 µL of each agonist (thrombin, U46619, ADP, convulxin) was added to the platelets, with final concentrations ranging from 0.1 to 20 µM, depending on the agonist. Platelet stimulation was carried out at room temperature for 10 minutes. After stimulation, 1 µL of FITC-conjugated anti-CD62P antibody (clone RB40.34, BD Biosciences) was added to each sample, and the platelets were incubated in the dark for an additional 10 minutes to prevent photobleaching.

Following staining, 100 µL of 2% formalin was added to fix the platelets, and samples were either immediately analyzed by flow cytometry or stored at 4°C for up to 16 hours prior to analysis. Flow cytometry was performed on a BD Accuri C6 cytometer, and data were collected for 10,000 events per sample. The expression of P-selectin (CD62P) on the platelet surface was used as an indicator of platelet activation, and data analysis was carried out using FlowJo software.

### Immunophenotyping by flow cytometry

At the designated time points, tumor-bearing left hemispheres were enzymatically digested using collagenase IV (Sigma) and DNase I (Sigma) to generate a single-cell suspension, subjected to a Percoll centrifugation step, and then filtered through a 40-μm strainer. Cells were first stained with LIVE/DEAD Fixable Stains (Thermo Fisher) on ice for 15 minutes, washed with PBS, and resuspended in an Fc receptor blocker (Miltenyi Biotec) diluted in PBS/2% BSA, followed by incubation on ice for 10 minutes. Surface staining was performed using fluorochrome-conjugated antibodies diluted in brilliant buffer (BD Biosciences) at a concentration of 1:100 to 1:250, with a 30-minute incubation on ice. After a subsequent wash in PBS/2% BSA, cells were fixed overnight using Foxp3/Transcription Factor Fixation Buffer (eBioscience).

For intracellular staining, antibodies were prepared in Foxp3/Transcription Factor Permeabilization Buffer at 1:250 to 1:500 and incubated at room temperature for 45 minutes. To detect intracellular cytokines, cells were stimulated in complete RPMI medium using a Cell Stimulation Cocktail with a protein transport inhibitor (eBioscience) for 4 hours. Following stimulation, cells underwent the same staining procedures as described above.

### In vitro CD8+ T cell culture

CD8+ T cells were extracted from the splenocytes of C57BL/6 mice using a magnetic bead isolation kit (STEMCELL Technologies), following established protocols. The cells were then maintained in complete RPMI medium supplemented with recombinant human IL-2 (30 U/mL, PeproTech) and activated using anti-CD3/anti-CD28 Dynabeads (Invitrogen) at a bead-to-cell ratio of 1:10. In addition, platelets were isolated as previously described and incorporated into the T cell culture for three consecutive days at a 1:1 ratio of CD8+ T cells to platelets. Cells subsequently underwent the same staining procedures outlined above.

### MEG-01 Cell Culture Experiments

MEG-01 cells were cultured in RPMI compete media as described above and treated with β-estradiol (0, 0.01, 0.1, 1.0, and 10 µg/ml), dihydrotestosterone (0, 0.01, 0.1, 1.0, and 10 µg/ml), and progesterone (0, 0.01, 0.1, 1.0, and 10 µg/ml) for 4 consecutive days for quantitative real time PCR (QPCR), immunoblotting and GPCR phospho antibody array (Full Moon Biosystems).

### Immunoblotting, Antibodies, and GPCR Phospho antibody array

Lysates were prepared as previousl described by our group^4, 45^. Cells were washed with PBS and lysed in NP-40 lysis buffer (0.5% NP-40, 10 mM Tris-Cl pH 7.5, 1 mM EDTA pH 8.0, 150 mM NaCl) supplemented with 1% protease and phosphatase inhibitors (Sigma-Aldrich). Lysates were clarified by centrifugation at 21,000 × g for 10 min at 4 °C, and protein concentration was determined using the Bio-Rad protein assay with BSA as the standard.

For immunoblotting, equal amounts of protein (15–50 µg) were resolved by SDS-PAGE on pre-cast 4–15% gels (Bio-Rad) and transferred to PVDF membranes (Millipore). The membrane was subsequently rinsed in Tris-buffered saline (TBS; 50 mM Tris-HCl 150 mM NaCl, pH 7.6) and blocked in Intercept blocking buffer for one hour (Li-Cor, 927-50000, Lincoln, NE) prior to the application of primary antibodies targeting Actin (1:1000) (Millipore Sigma, MAB1501PAR4, PAR4 (1:500) (Generated by Marvin Neiemen)^46^, SRC(Tyr418) (1:1000) (ThermoFisher, 14-9034-82), SRC (1:1000) (Cell Signaling 2108), P-PYK2(Tyr881) (1:1000) (ThermoFisher, 44-620G), and PYK2 (1:1000) (ABCAM, ab32571). The membrane was incubated overnight at 4°C with the primary antibody, rinsed with TBS-T (TBS; 50 mM Tris-HCl 150 mM NaCl, 1 % Tween-20 v/v% pH 7.6) followed by incubation in secondary (IRDye 680- and IRDye 800CW-conjugated secondary antibodies, Li-Cor, 926-68072 and 926-32213, respectively) for one hour at room temperature prior to visualization on the Li-Cor Odyssey CLx imaging System (Li-Cor, Lincoln, NE). Western blot analysis of band intensity was completed using Image Studio Analysis software from Li-Cor. Alternatively, Membranes were incubated with primary antibodies in 5% BSA in TBST at 4 °C for 2–7 days, washed with TBST, and then incubated with HRP-conjugated secondary antibodies for 2.5–3 h at room temperature. Blots were developed using enhanced chemiluminescence (ECL) and imaged on a Bio-Rad ChemiDoc MP system. Actin was used as a loading control.

For the GPCR Phospho antibody array (Full Moon Biosystems, PGP193) was conducted following the protocol as described by the company. Following array incubation in antibody, slides were sent to Full Moon Biosystems for analysis and array signal intensity was quantified as the median signal per spot and averaged across replicate spots for each antibody, with variability assessed by coefficient of variation. Data were normalized to the median signal intensity across the array. Fold changes were calculated as the ratio of treatment to control from normalized values, with increased or decreased expression indicated by red or green shading, respectively. For phospho-arrays, signaling ratios were calculated as the phospho-specific signal divided by the corresponding total protein signal, and fold changes were similarly determined between treatment and control samples.

### Co-immunoprecipitation

Co-Ips were done as previously described^47^. Briefly, Cells were lysed on ice for 2 h in 1.0 mL immunoprecipitation (IP) buffer containing 25 mM Tris (pH 7.4), 1 mM MgCl₂, 2 mM EDTA, 150 mM NaCl, 1.0% Triton X-100, and Complete Protease Inhibitor Cocktail (Roche). Protein G–Sepharose Fast Flow beads (100 μL; MilliporeSigma) were washed twice with IP buffer (500 × g, 5 min), blocked for 1 h in 5.0% BSA prepared in IP buffer, and resuspended in 1.0 mL IP buffer. Lysates were clarified by centrifugation (12,000 × g, 10 min, 4°C), and 10% of the supernatant (50 μL) was retained as the input control. The remaining lysate was split into two equal aliquots and incubated overnight at 4°C with Protein G beads and either 2 μg of the indicated primary antibody or an isotype-matched IgG control. Following incubation, samples were centrifuged at 500 × g for 5 min, and 50 μL of the supernatant was collected as the unbound fraction. Beads were washed four times with IP wash buffer (25 mM Tris [pH 7.4], 10 mM MgCl₂, 2 mM EDTA, 0.1% Triton X-100), with 50 μL aliquots retained after the first and fourth washes. After the final wash, bound proteins were eluted by denaturation in 2× SDS sample buffer and heating at 95°C for 10 min. Samples were resolved by SDS-PAGE and transferred to nitrocellulose membranes as described above.

### Quantitative real time PCR (qPCR)

MEG-01 cell were incubated at *37 °C in a humidified incubator with 5% CO₂, then rinsed with HBSS. RNA was isolated using the RNeasy Mini Kit, quantified for quality and concentration, and reverse-transcribed to cDNA. Quantitative PCR was performed using SYBR Green chemistry on a Bio-Rad CFX Connect system to assess expression of GSC, platelet, and coagulation-associated genes. Reactions (20 µL) were run in triplicate for 40 cycles, and relative expression was calculated using the ΔΔCT method.* Primers used were specific for actin (5’-AGAAAATCTGGCACCACACC-3’, 5’-AGAGGCGTACAGGGATAGCA-3’) and PAR4 (5’-CCGCTGCTGTATCCTTTGGT -3’, 5’-TCCTTGAGTTCTACTGTGGGAC-3’).

### MEG-01 calcium imaging

Washed MEG-01 were pre-incubated with BMS986120 (10nM), β-estradiol (0, 0.01, 0.1, 1.0, and 10 µg/ml), dihydrotestosterone (0, 0.01, 0.1, 1.0, and 10 µg/ml), and progesterone (0, 0.01, 0.1, 1.0, and 10 µg/ml). Afterwards, washed MEG-01 were loaded with Fura-2 AM (5 µM, 1 h at 37 °C), pelleted by centrifugation (2700 rpm, 5 min) in the presence of 10 µM PgI₂, and resuspended in fresh Tyrode’s solution in a 96-well plate. Intracellular calcium flux was measured over 10 min by ratiometric fluorescence (340/380 nm) following automated addition of thrombin (1.0 U/mL) using a FlexStation 3 system. Data are presented as normalized fluorescence changes relative to baseline (F/F₀) as previously described^43^.

### Analysis of previously published scRNA-seq data

#### Patient samples and datasets

This study utilized WHO grade 4 IDH wildtype GBM samples from four publicly available datasets: The Cancer Genome Atlas (TCGA), Chinese Glioma Genome Atlas (CGGA_325, CGGA_893), and Glioma Longitudinal Analysis (GLASS).

#### Immune Cell Profiling

The relative abundance of 22 immune cell subtypes within each tumor sample from each dataset was estimated using CIBERSORTx with default parameters and the LM22 signature gene set. Additionally, a gene signature for platelet expression (REACTOME_RESPONSE_TO_ELEVATED_PLATELET_CYTOSOLIC_CA2, MSigDB) was scored for each sample using single sample gene set enrichment analysis (ssGSEA).

#### Univariate survival analysis

To evaluate the prognostic significance of immune cell infiltration, univariate Cox proportional hazards models were constructed for each estimated immune cell population (including the Platelet Sig score) within each dataset. Immune cell proportions and platelet signature score were z-score normalized and centered prior to hazard ratio computation. Immune cell abundance and platelet signature score (platelet_sig) were dichotomized at the median value for each cell type within each respective dataset. The resulting hazard ratios (HRs) for overall survival are presented, with error bars indicating the 95% confidence intervals.

#### Immune cell correlation analysis

Spearman correlation coefficients between the estimated abundances of the major immune cell types and platelet gene signature score were computed. Correlation coefficients were computed for each of the major immune cell subtypes (B Cells, dendritic cells (DCs), T cells, NK cells, mast cells, microglia, monocytes, and macrophages) within each dataset (CGGA_325, GLASS, TCGA).

#### PAR4 expression analysis in cell subtypes

To investigate the expression profile of the PAR4 gene (also known as *F2RL3*) across different cell populations within the GBM TME, we analyzed single-cell RNA sequencing data from the GBMap dataset. The expression level of PAR4 was assessed for various identified cell subtypes as previously annotated. A dot plot (Seurat v5) was generated to visualize the results.

#### Mendelian randomization analysis

The European Ancestry consortium was used for this analysis. Genetic variants (G), typically single-nucleotide polymorphisms (SNPs), serve as instrumental variables (IVs) to assess the causal effect of an exposure (X; e.g., platelet count) on an outcome (Y; e.g., glioma risk). For the causal inference to be valid, three key assumptions must be satisfied: IV1 (Relevance) — the genetic instruments must be robustly associated with the exposure, typically derived from genome-wide association studies (GWAS); IV2 (Independence) — the genetic instruments must not be associated with confounding variables (U) that jointly influence both the exposure and the outcome; and IV3 (Exclusion Restriction) — the instruments must affect the outcome solely through the exposure and not via any alternative biological pathways. The directed acyclic graph visualizes how genetic variants influence the outcome only via their effect on the exposure, assuming no horizontal pleiotropy or confounding from unmeasured variables.

#### Akoya Multispectral Immunofluorescence

##### PhenoCycler-Fusion Slide Preparation

FFPE tissue slides were baked at 60 °C overnight. Baked slides were deparaffinized with Xylene Substitute solution (Epredia) and rehydrated in a series of gradient ethanol solutions, ranging from 100% to ddH2O. The slides were then placed in a glass Coplin jar containing 1X antigen retrieval buffer (pH 9.0) (Akoya Biosciences), and antigen retrieval was performed with an Instant Pot using the high-pressure setting for 20 minutes. Following antigen retrieval, the slides were cooled in the retrieval buffer to RT and transferred to a transparent slide container with an autofluorescence bleaching solution (4.5% w/v H2O2 and 20mM NaOH in PBS). The container was sandwiched between two LED lamps, and the slides were photobleached for 90 minutes at RT, with fresh bleaching solution replenished once after 45 minutes. Slides were then processed according to the PhenoCycler-Fusion User Guide (PD-000011 Rev S, Akoya Biosciences). Briefly, slides were washed in Hydration Buffer, equilibrated in Staining Buffer, and incubated overnight at 4 °C with a 47-marker antibody cocktail (Table S8) prepared in Blocking Buffer. The slides were then washed in Staining Buffer, gently fixed with 1.6% formaldehyde, washed in PBS, and incubated in ice-cold methanol for 5 minutes. The slides were then washed in PBS, fixed with Final Fixative Solution for 20 minutes, washed in PBS, and preserved in Storage Buffer at 4 °C before the PCF run.

#### PhenoCycler-Fusion Setup and Data Acquisition

The experimental protocol was set up using the PhenoCycler Experiment Designer (Version 2.2.0). A reporter plate containing fluorescently labeled reporters was prepared according to the PhenoCycler-Fusion User Guide (PD-000011 Rev S, Akoya Biosciences). Slides were prepared for the PCF instrument according to the PhenoImager Fusion 2.0 User Guide (PD-000088 Rev G, Akoya Biosciences). Briefly, slides were removed from the Storage Buffer and incubated in PBS for 10 minutes at RT. After incubation, a Flow Cell (Akoya Biosciences) was attached to each sample slide using the Flow Cell Assembly Device (Akoya Biosciences). The slide with the attached Flow Cell (Sample Flow Cell) was equilibrated in 1X PCF buffer for 10 minutes at RT. PhenoCycler-Fusion software (Version 2.4.0) was used to set up the imaging run on the PCF instrument according to the PhenoImager Fusion 2.0 User Guide (PD-000088 Rev G, Akoya Biosciences).

Reagents were prepared and loaded into appropriate reservoirs on the instrument, and the prepared reporter plate was loaded into the instrument. A new PhenoCycler run was initiated using the experimental protocol design. A blank flow cell was loaded into the Flow Cell Slide Carrier, and all software prompts during the pre-flight routine were followed. The Sample Flow Cell was loaded into the carrier, and a leak check was performed. Automatic sample detection was performed, followed by scan region selection and imaging initiation. Upon completion, the Sample Flow Cell was placed in the Storage Buffer at 4 °C. The generated QPTIFF data files were used for downstream image analysis. Analysis of the akoya multispectral immunofluorescence imaging was done as previously described^48^.

#### PhenoCycler-Fusion data analysis

We leveraged InstanSeg^49^, an embedding-based deep learning model built for cell segmentation, to identify cell boundaries in each region of interest (ROI) extracted from the highly multiplexed immunofluorescence images generated by the Akoya PhenoCycler Fusion. The resulting segmentation mask and a list of de-multiplexed single-channel images from the panel are input to Nimbus^50^, a deep learning model that predicts confidence of marker positivity in each cell. These scores, calculated for each channel and bounded between 0 to 1, replace integrated expression for downstream analyses to combat technical confounders like background staining, autofluorescence, and channel intensity variation.

By aggregating Nimbus scores across every high quality ROI into a single AnnData object, we applied FlowSOM^51^, an unsupervised clustering algorithm based on self-organizing maps, to group cells by their pattern of protein expression. Using the markers enriched in each of the 18 resulting clusters, as identified by the Wilcoxon rank sum test, we manually annotated cell states. We then appended these annotations to the spatial coordinates of each cell for further analysis.

To capture acellular features in each ROI, such as platelets, we employed a deep learning model for single-channel bright spot detection in fluorescence microscopy images called Spotiflow^52^. To efficiently calculate the density and proximity of these features around cells, we applied the cKDTree algorithm from the scipy.spatial library^53^. By aggregating these spatial association metrics for each previously defined cell state and comparing across ROI-specific metadata, we investigated sex-specific and treatment-correlated phenotypes.

#### Code and data availability

All data used for the above analyses are publicly available through the TCGA, CGGA, GLASS, and GBMap datasets, including clinical and cell type metadata. Immune cell signatures are available through CIBERSORTx, and the platelet gene signature is available through MSigDB. All analyses were performed in R version 4.3.0, and all code to reproducibly perform these analyses will be provided on Github.

#### Statistical analysis

Data visualization and statistical analysis were conducted using GraphPad Prism (Version 9, GraphPad Software Inc., RRID:SCR_002798). Depending on the dataset, unpaired or paired t-tests, as well as one-way or two-way analysis of variance (ANOVA) with multiple comparison adjustments, were applied as described in the figure legends. Survival analysis was evaluated using the log-rank test. For all analysis, statistical significance was set at p < 0.05.

## Data Availability

All data generated in this study are available upon request from the corresponding author, Dr. Justin D. Lathia.

## Code availability

All code to reproducibly perform these analyses will be provided on Github. Additionally, all code used in this study are available upon request from the corresponding author, Dr. Justin D. Lathia.

## Author Contributions

Conception and design: A.R.S, S.J.C, J.D.L Development of methodology: A.R.S, G.B, A.A, J.L

Acquisition of data: A.R.S, G.B, D.R, T.J.A, I.J, V.B, L.T, T.N, N.R, S.H, G.P.T, A.V, J.G, X.Y, S.K, J.C, V.S.K, S.S.K, A.S, B. R., N.S., A.D

Analysis and interpretation of data: A.R.S, G.B, J. L

Writing review: A.R.S, J.L, E.M.H, J.D.R, C.M.O, S.J.C, J.D.L

Administrative, technical, or material support: C.N.H, A.E.S, J.B.R, C.G.H, E.X.S, F.W.L, W.A.G, T.E.M, C.M.O, T.A.C, A.K, M.N., A.D.

Study supervision: S.J.C, J.D.L.

## Acknowledgements

We sincerely acknowledge the members of the Lathia laboratory for their valuable insights and constructive discussions and Dr. Reza Khatib for his inspiration and support of our work. We sincerely want to acknowledge Neal Restivo and Oatey Company for their support of our work. We are grateful for the illustrative contributions of Ms. Amanda Mendelsohn from Enterprise Creative at the Cleveland Clinic. We are grateful to Dr. Belinda Willard and her team at the Proteomics Core (Cleveland Clinic Research). Additionally, we extend our gratitude to Drs. Matthew Flick and Silvio Antoniak (University of North Carolina Chapel Hill) for their support and constructive comments on the project. This work was supported by NIH grants NIH T32 CA059366 (A.R.S), F32 CA287655 (A.R.S), R35 NS127083 (J.D.L), P01 CA245705 (J.D.L, J.B.R), HL158801 (SJC), U01HL143402 (AK), and R01HL164516 (A.K.). This work was also supported by the Cleveland Clinic VeloSano Bike to Cure event (S.J.C and J.D.L), Velosano 11 Pilot Grant (S.J.C), and the Midwest Brain Tumor Foundation (A.R.S), Cleveland Clinic (A.K., J.D.L.) and Case Comprehensive Cancer Center (J.D.L.).

