## Supplementary material for "Platelets regulate glioblastoma growth and immunity via sex-dependent PAR4 - Estrogen receptor beta signaling": SloanLathia_PAR4 Extended Data _ Clean.pdf

### Supplementary Materials

#### Supplementary Figures

Supplementary Figure 1. Platelet gene signature correlates to GBM progression

Supplementary Figure 2. GBM platelet reactivity compared to healthy subject platelets

Supplementary Figure 3. Drug-target Mendelian randomization (MR) analysis of PAR4 variants proxying lowered platelet count and their effects on glioma risk by age group

Supplementary Figure 4. Platelet depletion prolongs survival in a sex-dependent manner

Supplementary Figure 5. Inhibiting the thrombin-PAR4 signaling axis prolongs survival in females and is not PAR4 expression dependent

Supplementary Figure 6. *In vitro* MEG-01 PAR4 expression with hormone treatment

Supplementary Figure 7. Inhibiting the thrombin-PAR4 signaling axis prolongs survival in a sex-dependent manner through the TME

Supplementary Figure 8. Tumor-infiltrating T cell subsets with 14-day BMS986120 treatment

Supplementary Figure 9. Tumor-infiltrating CD8+ T cell number and function with 10-day BMS986120 treatment

Supplementary Figure 10. Tumor-infiltrating CD8+ T cell number and function with 7-day BMS986120 treatment

Supplementary Figure 11. Tumor-infiltrating non-T cell subsets with 14-day BMS986120 treatment

Supplementary Figure 12. Tumor-infiltrating myeloid subsets with 14-day BMS986120 treatment

Supplementary Figure 13. Tumor-infiltrating immune cell flow cytometry clean up

Supplementary Figure 14. Tumor-infiltrating myeloid subsets flow cytometry clean up

Supplementary Figure 15. BMS986120 improves survival in C57BL/6 FOXP3-DTR mice

Supplementary Figure 16. BMS986120 decreases intratumoral platelet expression and increase CD8+ T cells in females but not males

Supplementary Figure 17. PAR1 but not PAR4 is expressed on mouse T cells

Supplementary Figure 18. PAR1 but not PAR4 is expressed on human T cells

Supplementary Figure 19. Sex biases between male and female platelets exist regardless of BMS986120 administration

Supplementary Figure 20. Tumor-bearing platelet reactivity in immune-incompetent mice

#### Supplementary Data Tables

Supplementary Table 1. Platelet Gene Signature Genes

Supplementary Table 2. Platelet signature score

Supplementary Table 3. Demographics for matched control subjects

Supplementary Table 4. Demographics for GBM patients

Supplementary Table 5. Flow antibodies for immune cell subsets

Supplementary Table 6. Flow antibodies for cytokine profiling

Supplementary Table 7. Flow antibodies for myeloid immune cell subset profiling

**Extended Data**

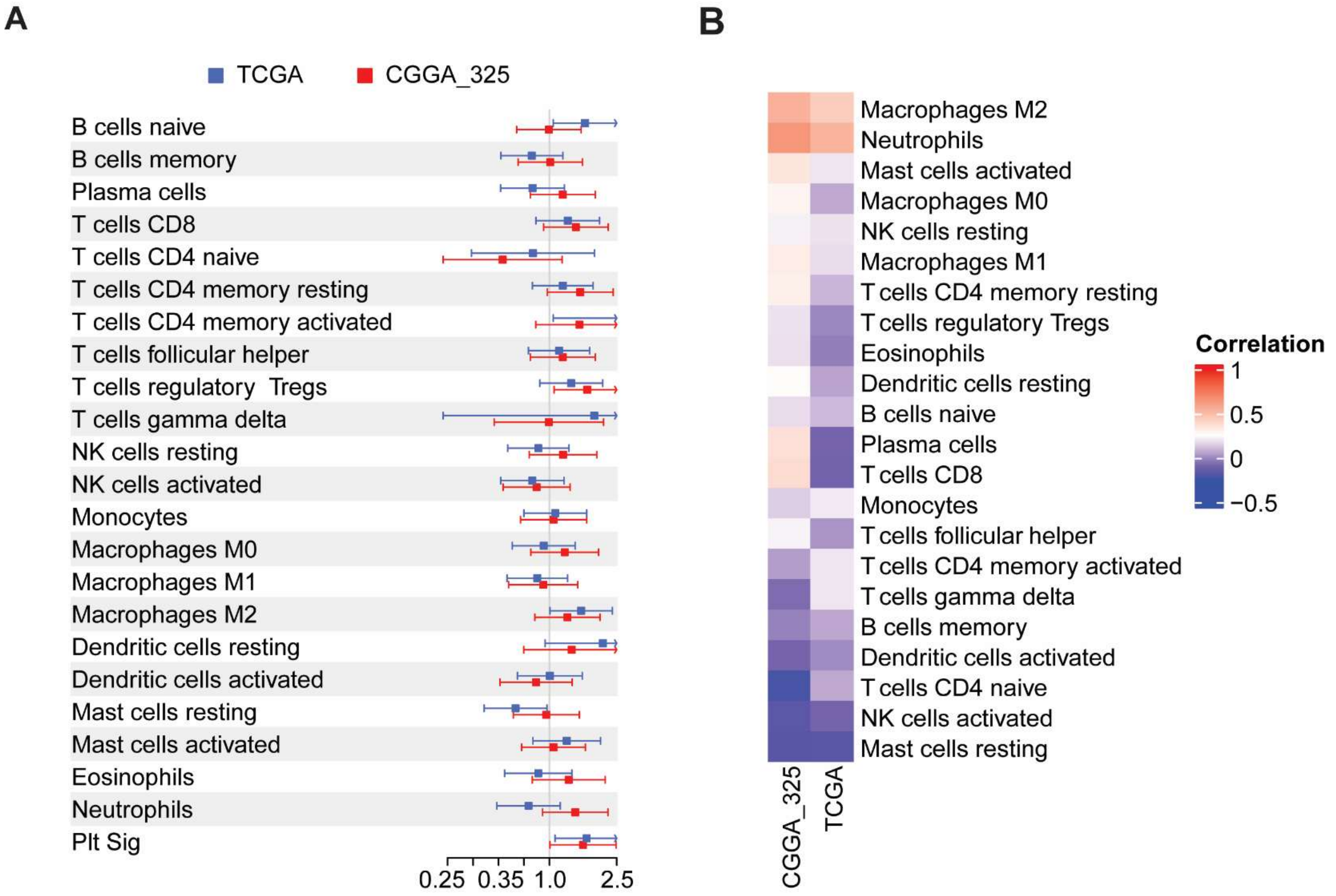


**Supplementary Figure 1. Platelet gene signature correlates with GBM progression. A**. WHO grade 4 IDH wildtype glioblastoma samples from The Cancer Genome Atlas (TCGA) and Chinese Glioma Genome Atlas (CGGA) publicly available data sets. Univariate Cox proportional hazards models were constructed, and immune cell proportions and platelet signature score were z-score normalized and centered prior to hazard ratio computation. **B**. WHO grade 4 IDH-wildtype GBM samples from The Cancer Genome Atlas (TCGA) and Chinese Glioma Genome Atlas (CGGA) publicly available data sets. Spearman correlation coefficients were computed between the estimated abundances of the major immune cell types and platelet gene signature score.

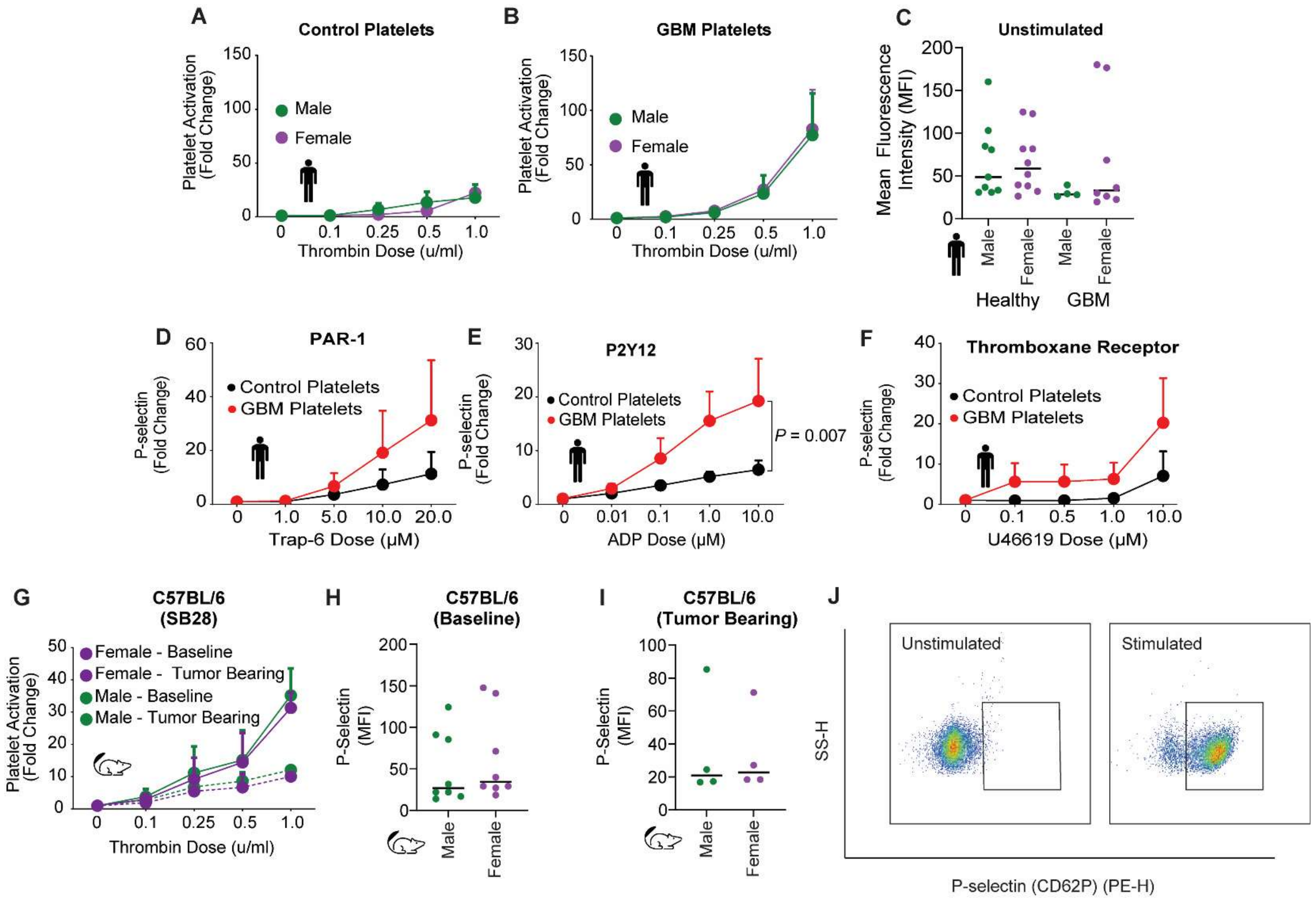


**Supplementary Figure 2. GBM platelet reactivity compared to control subject platelets. A.** Sex stratification of washed platelets isolated from matched control subjects (**A**) and patients with a GBM diagnosis (**B**). Platelets were treated with AY-NH2 to stimulate PAR4, and α-granule secretion was measured using an antibody specific for P-selectin via flow cytometry. **C**. Sex stratification of unstimulated washed platelets isolated from matched control subjects and patients with a GBM diagnosis. Spontaneous α-granule secretion was measured using an antibody specific for P-selectin via flow cytometry. Data are represented as means ± SEM from *n*=18 GBM patients and *n*=18 control subjects. Two-way ANOVA was performed. **D-F**. Washed platelets isolated from patients with a GBM diagnosis and matched control subjects were treated with TRAP-6 to stimulate PAR1 (**D**), ADP to stimulate P2Y12 (**E**), and U46619 to stimulate the thromboxane receptor (**F**), and α-granule secretion was measured using an antibody specific for P-selectin via flow cytometry. **G**. Sex stratification of washed platelets isolated from mice before intracranial tumor implantation (baseline) and following intracranial injection of the murine GBM SB28. Following tumor formation, washed platelets were isolated, and α-granule secretion was measured using an antibody specific for P-selectin via flow cytometry. Data are represented as means ± SEM from n=4 independent experiments. Two-way ANOVA was performed. **H**, **I**. Sex stratification of washed platelets isolated from mice before intracranial tumor implantation (baseline) (**H**) and following intracranial injection (**I**) of the murine GBM SB28. Spontaneous α-granule secretion was measured using an antibody specific for P-selectin via flow cytometry. Data are represented as means ± SEM from n=4 independent experiments (tumor-bearing) and n=8 animals for baseline. An unpaired student's *t*-test was performed. **J**. Gating strategy for measuring α-granule secretion via CD62P in unstimulated or agonist-induced platelets isolated from human or mouse.

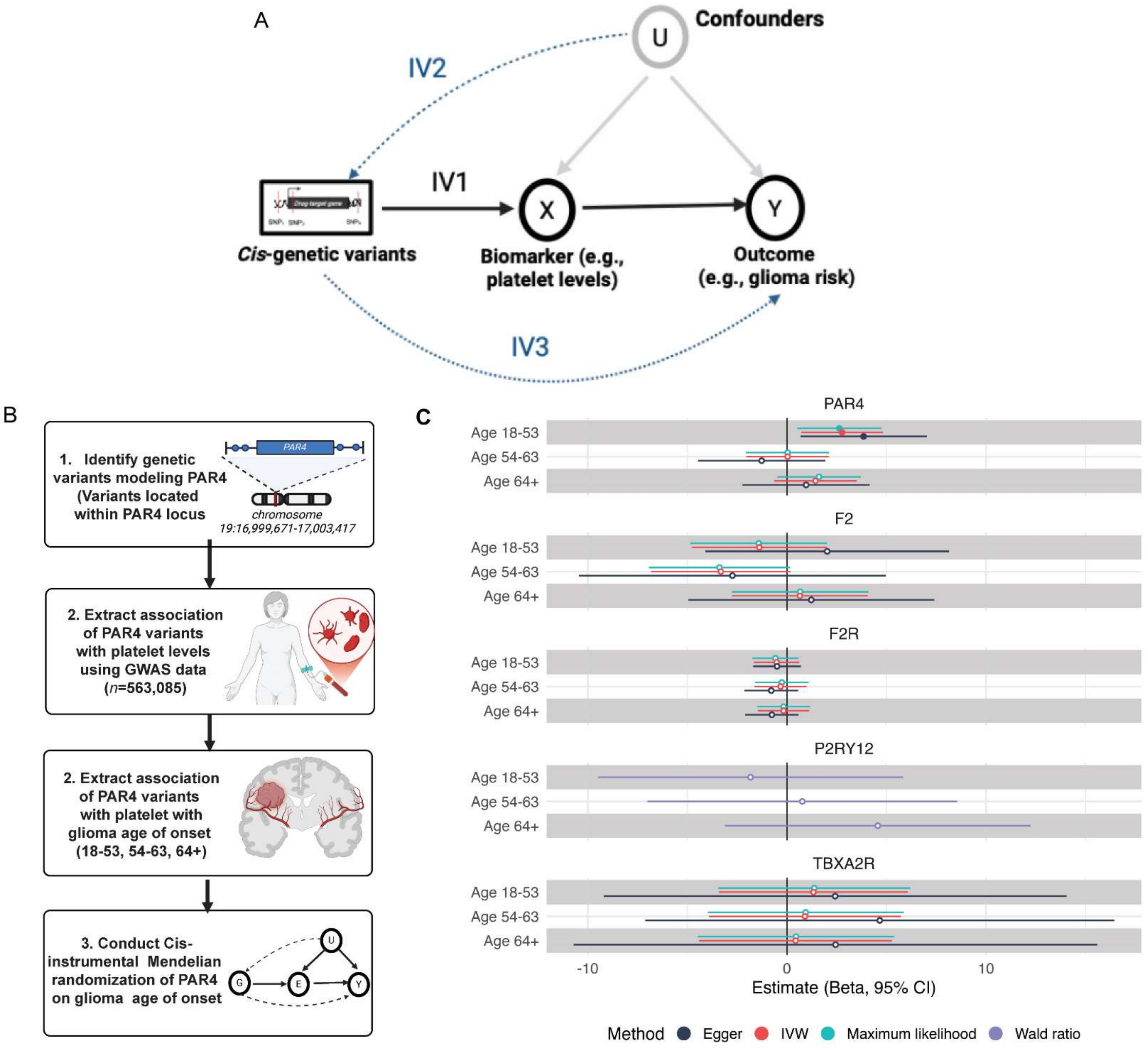


**Supplementary Figure 3. Drug-target Mendelian randomization (MR) analysis of PAR4 variants proxying lowered platelet count and their effects on glioma risk by age group. A.** This diagram illustrates the core assumptions underlying Mendelian randomization (MR). In this framework, genetic variants (G), typically single nucleotide polymorphisms (SNPs), serve as instrumental variables (IVs) to assess the causal effect of an exposure (X; e.g., platelet count) on an outcome (Y; e.g., glioma risk). For the causal inference to be valid, three key assumptions must be satisfied: IV1 (Relevance) — the genetic instruments must be robustly associated with the exposure, typically derived from genome-wide association studies (GWAS); IV2 (Independence) — the genetic instruments must not be associated with confounding variables (U) that jointly influence both the exposure and the outcome; and IV3 (Exclusion Restriction) — the instruments must affect the outcome solely through the exposure and not via any alternative biological pathways. The DAG visualizes how genetic variants influence the outcome only via their effect on the exposure, assuming no horizontal pleiotropy or confounding from unmeasured variables. **B.** Overview of the MR analysis framework: Variants in the *PAR4* locus were used as instrumental variables to proxy the effect of lowered platelet count on glioma risk. The analysis was stratified into three age-of-onset groups: 18-53 years, 54-63 years, and 64+ years. **C**. Forest plot of the main MR results: Inverse variance weighted (IVW) estimates are shown alongside sensitivity analyses, including MR-Egger regression and maximum likelihood approaches. Results are presented with 95% confidence intervals to assess the robustness of the findings.

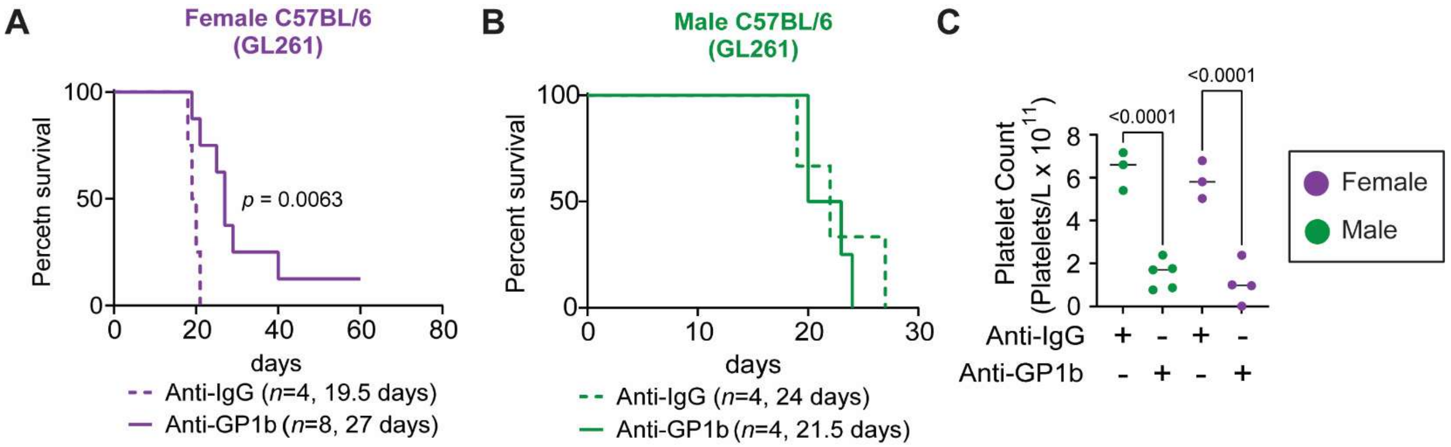


**Supplementary Figure 4. Platelet depletion prolongs survival in a sex-dependent manner.** Kaplan-Meier survival analysis was performed after intracranial tumor implantation of mouse GBM cell model GL261 in C57BL/6 female (**A**) and male (**B**) mice treated with anti-GP1bα depleting antibody. Statistical significance for survival analysis was determined by log-rank test. **C**. Complete blood count (CBC) platelet reading 1 day post anti-GP1bα to validate platelet depletion in tumor-bearing mice used in panels a and b. Data are shown with each point representing an individual animal. One-way ANOVA analysis with Tukey's multiple comparison test was performed.

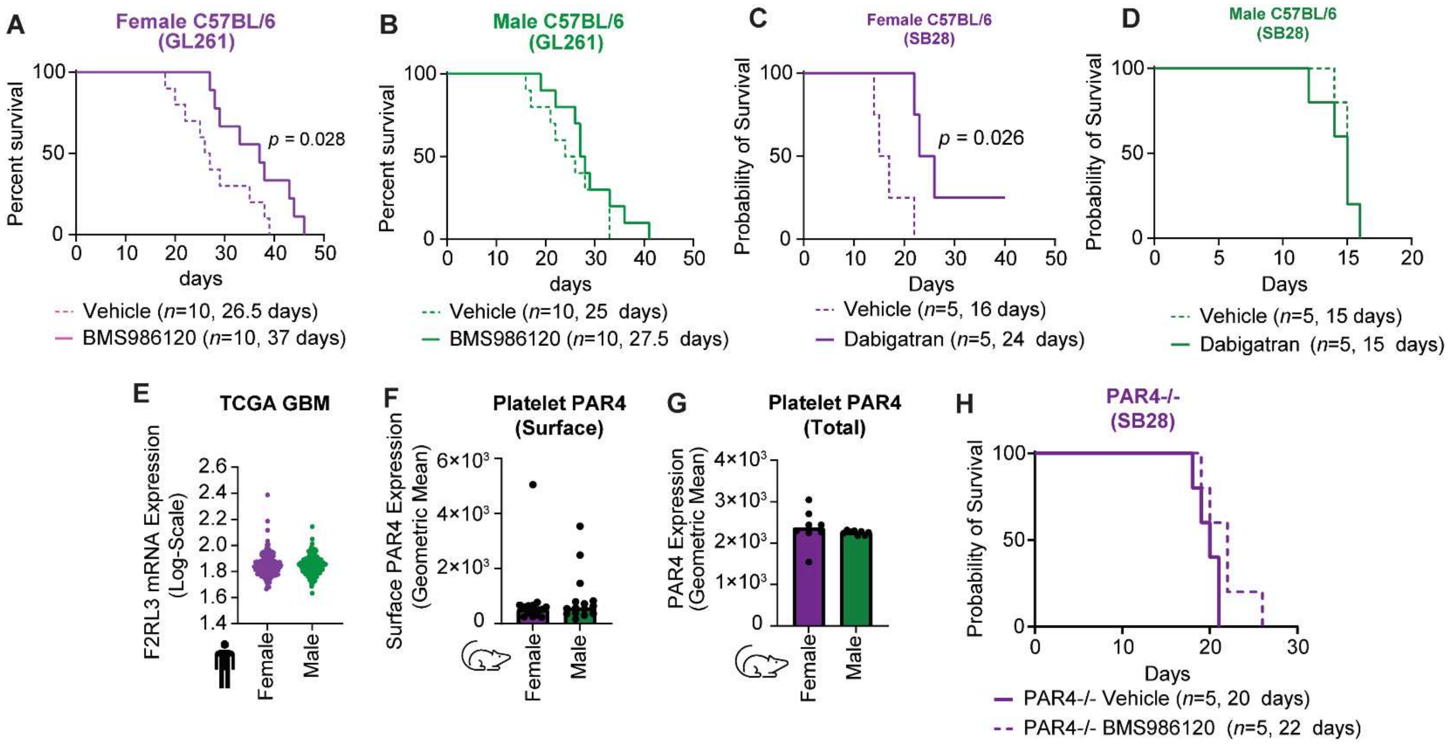


**Supplementary Figure 5. Inhibiting the thrombin-PAR4 signaling axis prolongs survival in females and is not dependent on PAR4 expression. A, B.** Kaplan-Meier survival analysis was performed after intracranial implantation of the mouse GBM cell line GL261 in immunocompetent C57BL/6 mice followed by pharmacologically inhibiting the thrombin PAR4 signaling axis by administering BMS986120 (2 mg/kg), a selective PAR4 inhibitor. Statistical significance for survival analysis was determined by log-rank test. **C, D.** Kaplan-Meier survival analysis was performed after intracranial implantation of the mouse GBM cell line SB28 in immunocompetent C57BL/6 mice followed by pharmacologically inhibiting the thrombin PAR4 signaling axis by administering dabigatran (50 mg/kg), a direct thrombin inhibitor. Statistical significance for survival analysis was determined by log-rank test. **E**. TCGA analysis of PAR4 gene expression in male and female tumors. **F**. Surface PAR4 expression measured by flow cytometry in female and male C57BL/6 unmanipulated animals. **G**. Total PAR4 expression measured by flow cytometry following membrane permeabilization in female and male unmanipulated animals. Data are shown with each point representing an individual animal. One-way ANOVA analysis with Tukey's multiple comparison test was performed. **H**. Kaplan-Meier survival analysis was performed after intracranial implantation of the mouse GBM cell line SB28 in PAR4-/- mice followed by pharmacologically inhibiting the thrombin PAR4 signaling axis by administering BMS986120 (2 mg/kg), a selective PAR4 inhibitor. Statistical significance for survival analysis was determined by log-rank test.

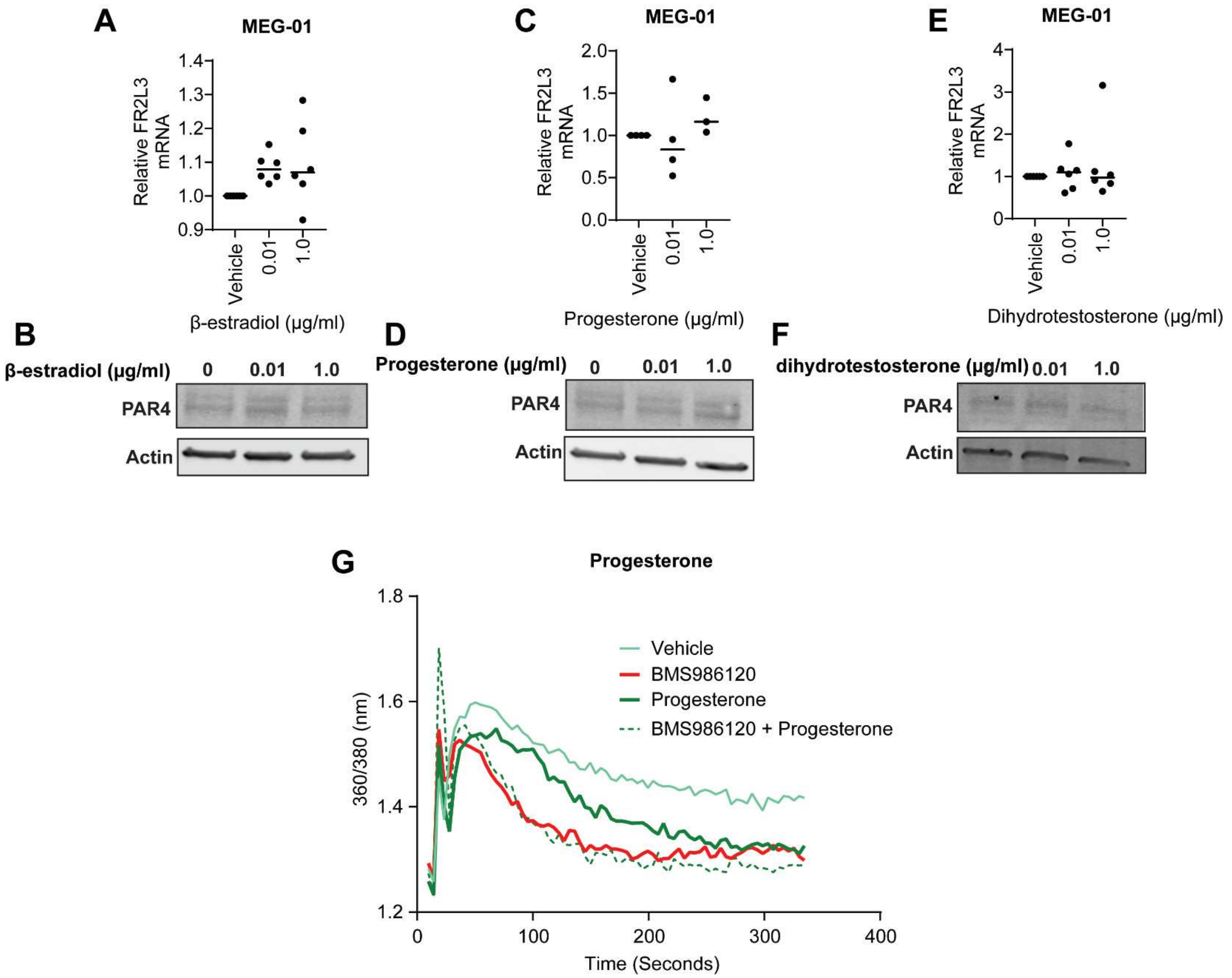


**Supplementary Figure 6.** ***In vitro*** **MEG-01 PAR4 expression with hormone treatment.** MEG-01 pre-platelet cells were treated with β-estradiol (**A**, **B**), progesterone (**C**,**D**), or dihydrotestosterone (**E**,**F**) for 4 consecutive days, and *F2rl3* (PAR4-encoding gene) mRNA expression (**A**, **C**, **E**) and PAR4 protein expression (**B**, **D**, **F**) were measured. **G**. Fura-2 calcium imaging on MEG-01 cells pretreated with BMS986120 and/or progesterone followed by stimulation with thrombin.

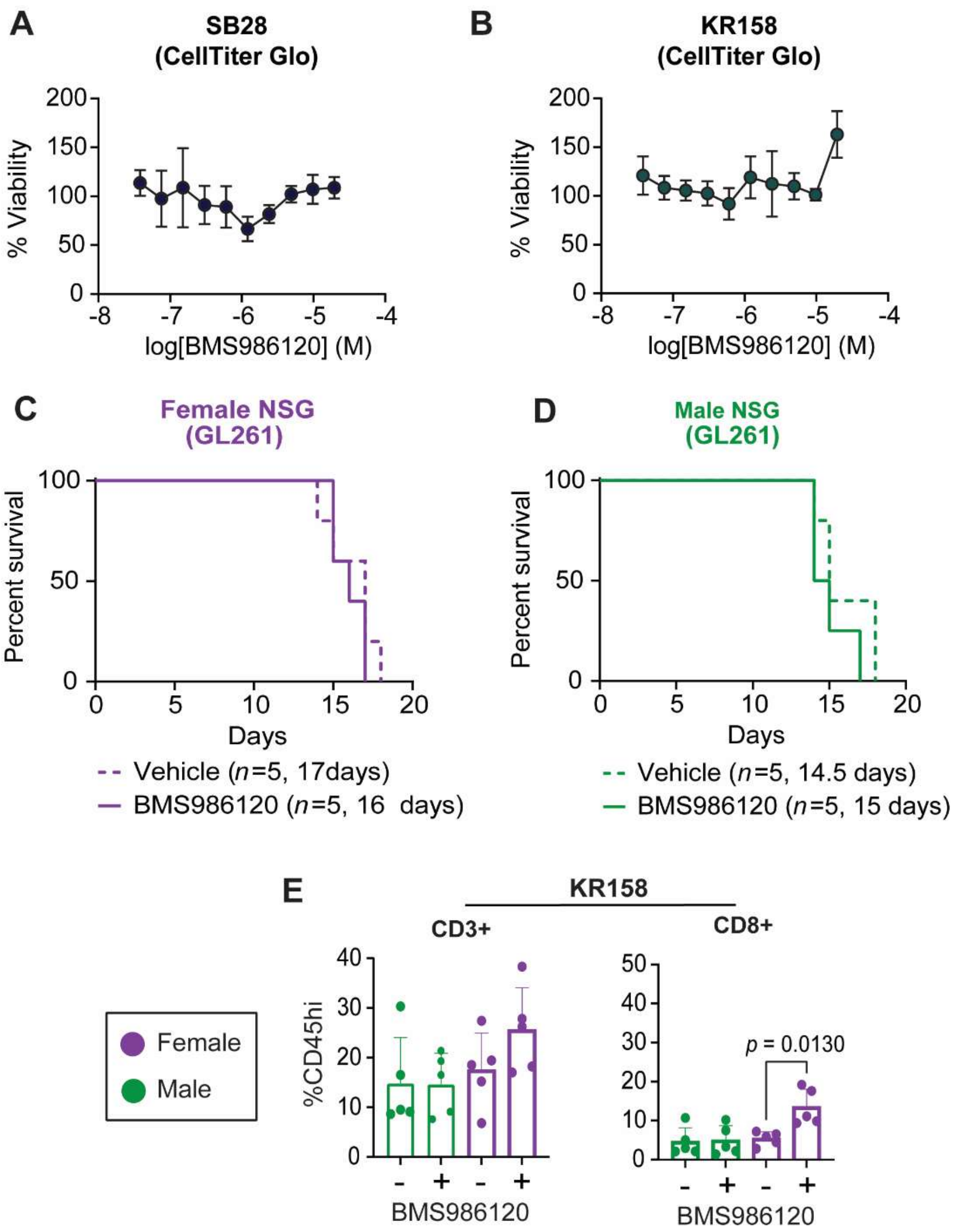


**Supplementary Figure 7. Inhibiting the thrombin-PAR4 signaling axis prolongs survival in a sex-dependent manner through the TME. A, B**. $IC_{50}$ analysis of mouse GBM cell lines (**A**) SB28 and (**B**) KR158 treated with escalating doses of BMS986120. **C, D**. Kaplan-Meier survival analysis was performed after intracranial implantation of the mouse GBM cell line GL261 in immunodeficient NSG mice followed by pharmacologically inhibiting the thrombin-PAR4 signaling axis by administering BMS986120 (2 mg/kg). Statistical significance for survival analysis was determined by log-rank test. **E**. Frequency of tumor-infiltrating T cell subsets 35 days post intracranial implantation of KR158 in male and female C57BL/6 mice treated with and without BMS98612. Data are combined from two independent experiments. One-way ANOVA analysis with Tukey's multiple comparison test was performed.

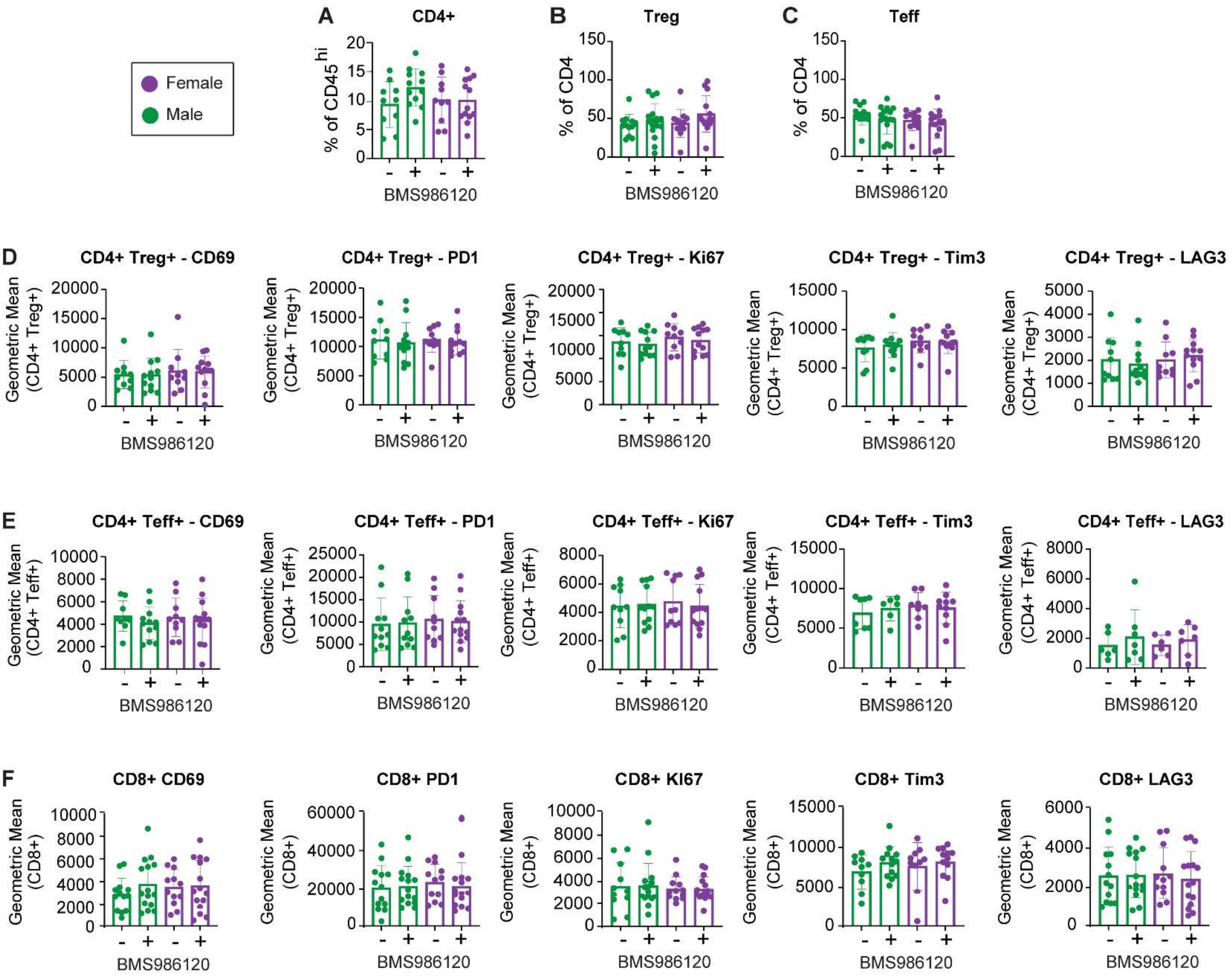


**Supplementary Figure 8. Tumor-infiltrating T cell subsets with 14-day BMS986120 treatment.** Frequency of tumor-infiltrating T cell subsets 14 days post intracranial implantation of SB28 cells in male and female C57BL/6 mice treated with and without BMS986120. **A-C**. Frequency of CD4+ T cell subsets. **D**. Frequency of tumor-infiltrating CD4+ Treg proliferation, exhaustion, and inhibitory receptor markers. **E**. Frequency of tumor-infiltrating CD4+ Teff proliferation, exhaustion, and inhibitory receptor markers. **F**. Frequency of CD8+ proliferation, exhaustion, and inhibitory receptor markers. Data are combined from two independent experiments.

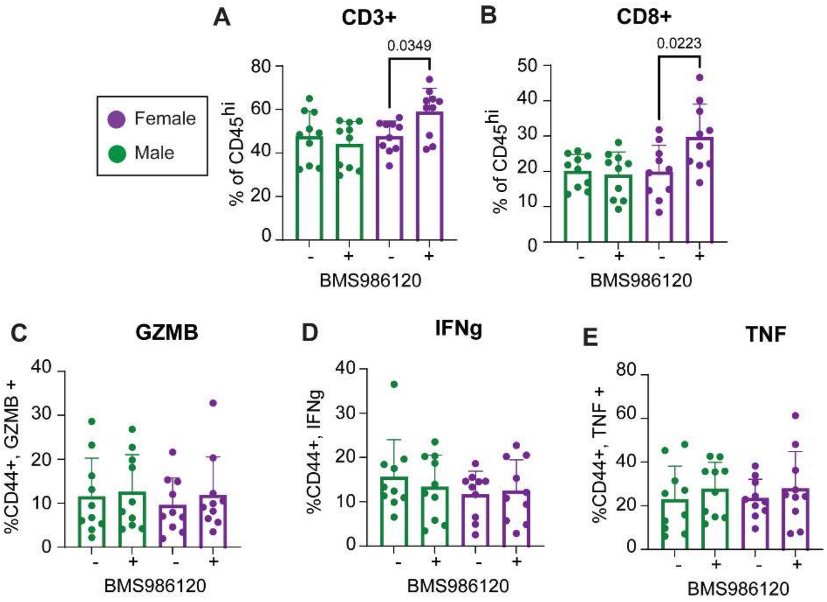


**Supplementary Figure 9. Tumor-infiltrating CD8+ T cell number and function with 10-day BMS986120 treatment.** Frequency of tumor-infiltrating T cell subsets 10 days post intracranial implantation of SB28 cells in male and female C57BL/6 mice treated with and without BMS986120. **A**. Frequency of CD3+ T cell subsets. **B**. Frequency of tumor-infiltrating CD8+ T cell subsets. **C-E**. Intracellular cytokine expression in tumor-infiltrating CD8+ T cells with 10 days BMS986120 treatment. Data are combined from two independent experiments. One-way ANOVA analysis with Tukey's multiple comparison test was performed.

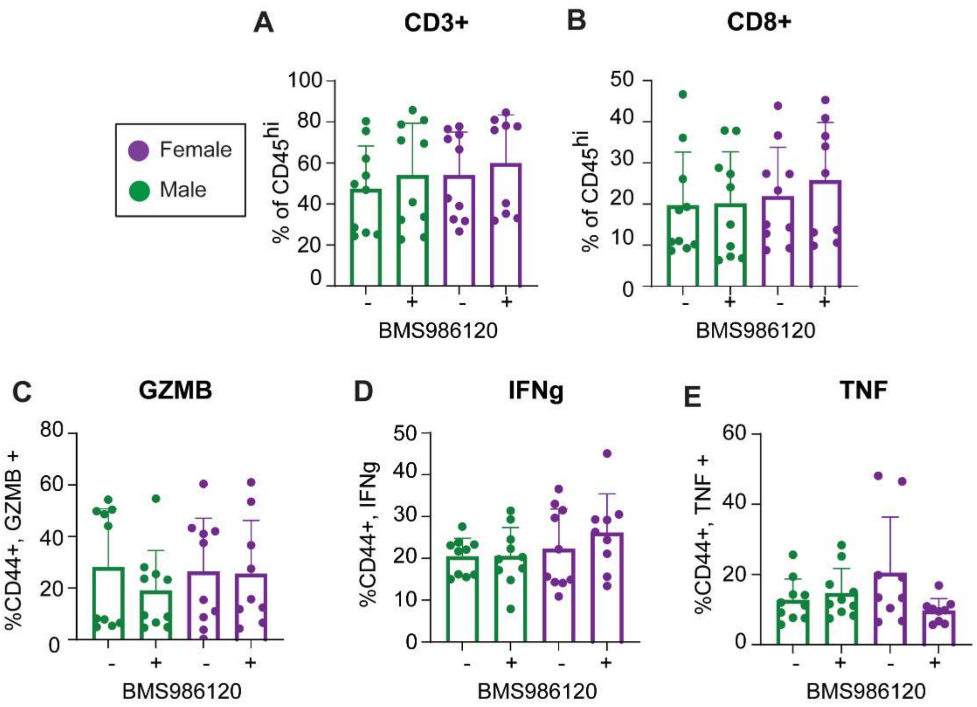


**Supplementary Figure 10. Tumor-infiltrating T cell subsets with 7-day BMS986120 treatment.** Frequency of tumor-infiltrating T cell subsets 10 days post intracranial implantation of SB28 cells in male and female C57BL/6 mice treated with and without BMS986120. **A**. Frequency of CD3+ T cell subsets. **B**. Frequency of tumor-infiltrating CD8+ T cell subsets. **C-E**. Intracellular cytokine expression in tumor-infiltrating CD8+ T cells with 10 days BMS986120 treatment. Data are combined from two independent experiments. One-way ANOVA analysis with Tukey's multiple comparison test was performed.

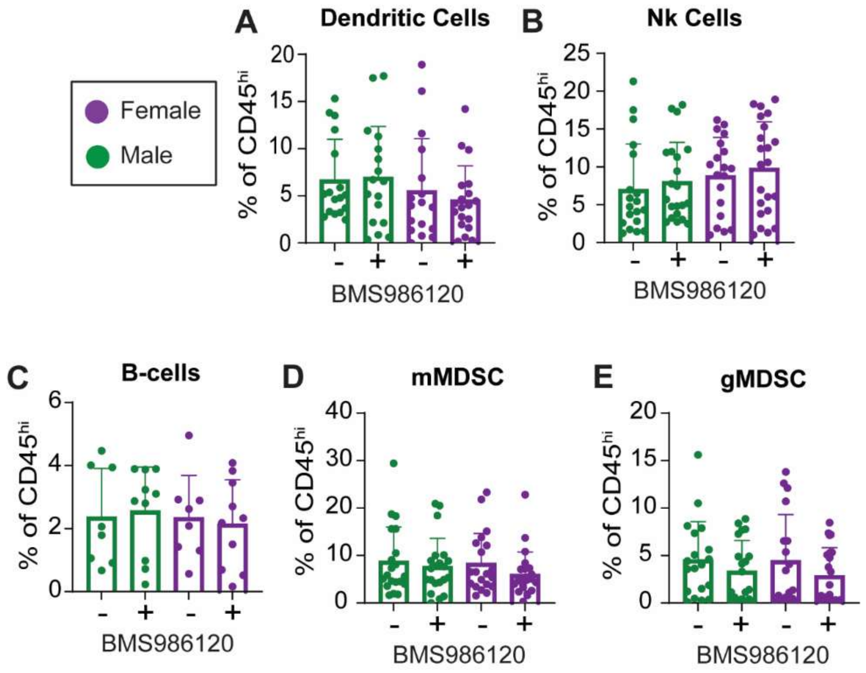


**Supplementary Figure 11. Tumor-infiltrating non-T cell subsets with 14-day BMS986120 treatment.** Frequency of tumor-infiltrating T cell subsets 14 days post intracranial implantation of SB28 cells in male and female C57BL/6 mice treated with and without BMS986120. Frequency of (**A**) dendritic cells, (**B**) NK cells, (**C**) B cells, (**D**) gMDSCs, and (**E**) mMDSCs. Data are combined from two independent experiments. One-way ANOVA analysis with Tukey’s multiple comparison test was performed.

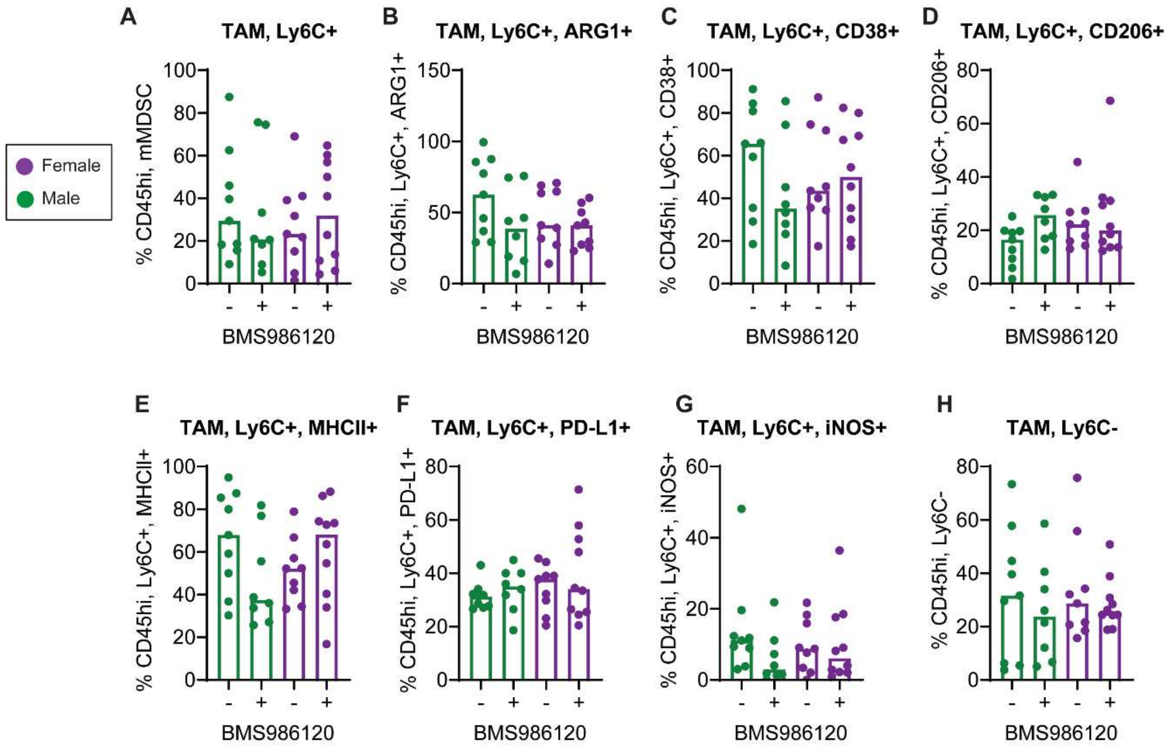


**Supplementary Figure 12. Tumor-infiltrating myeloid subsets with 14-day BMS986120 treatment.** Frequency of tumor-infiltrating T cell subsets 14 days post intracranial implantation of SB28 cells in male and female C57BL/6 mice treated with and without BMS986120. Frequency of (**A**) TAM, Ly6C+, (**B**) TAM, Ly6C+, ARG1+, (**C**) TAM, Ly6C+, CD38+, (**D**) TAM, Ly6C+, CD206+, (**E**) TAM, Ly6C+, MHCII+, (**F**) TAM, Ly6C+, PD-L1+, (**G**) TAM, Ly6C+, iNOS+, and (**H**) TAM, Ly6C-. Data are combined from two independent experiments. One-way ANOVA analysis with Tukey’s multiple comparison test was performed.

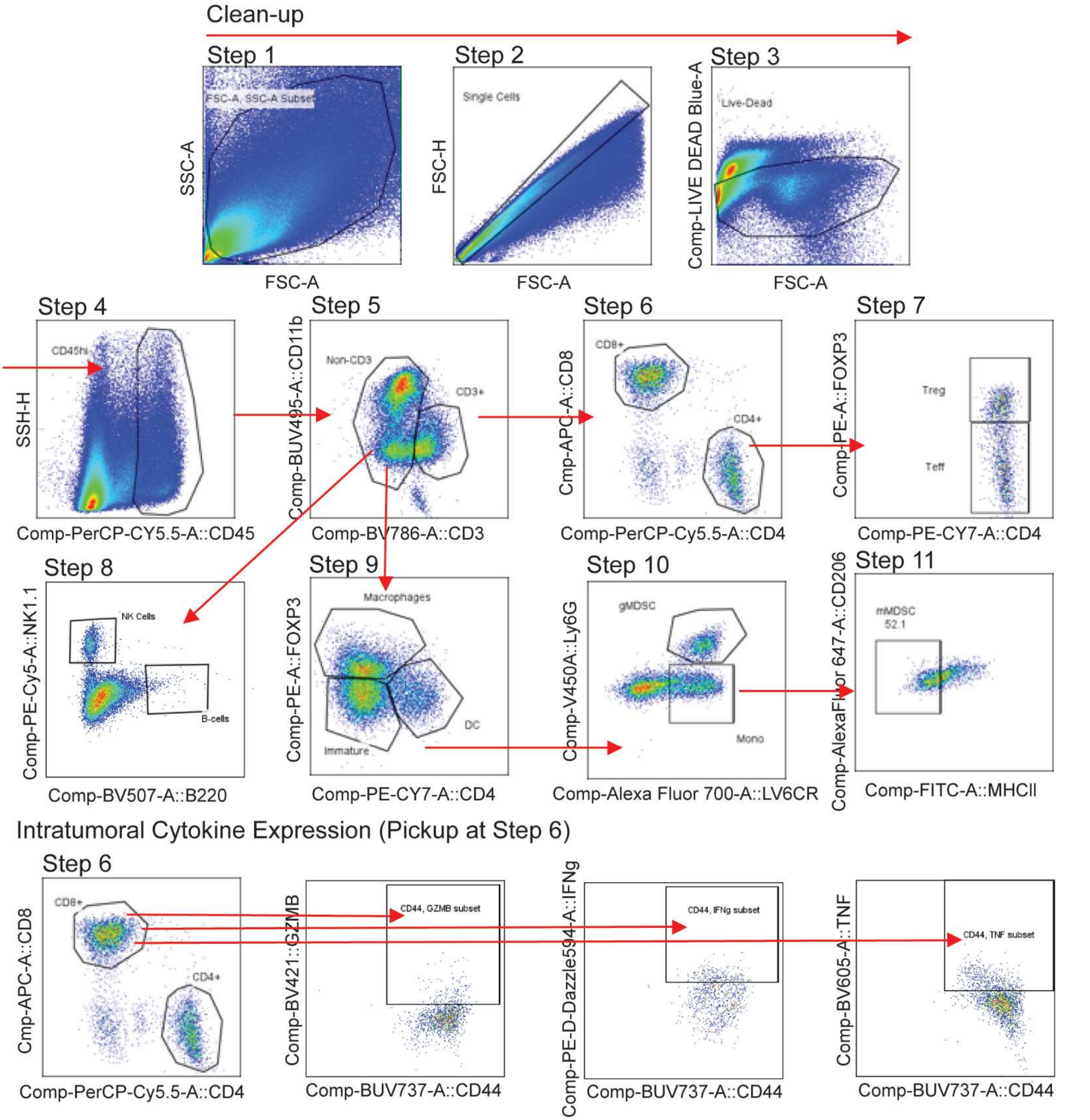


**Supplementary Figure 13. Tumor-infiltrating immune cell flow cytometry clean up.** Gating strategy for the analysis of tumor-infiltrating immune cell populations and CD8+ cytokine expression in male and female mice treated with BMS986120 relative to vehicle.

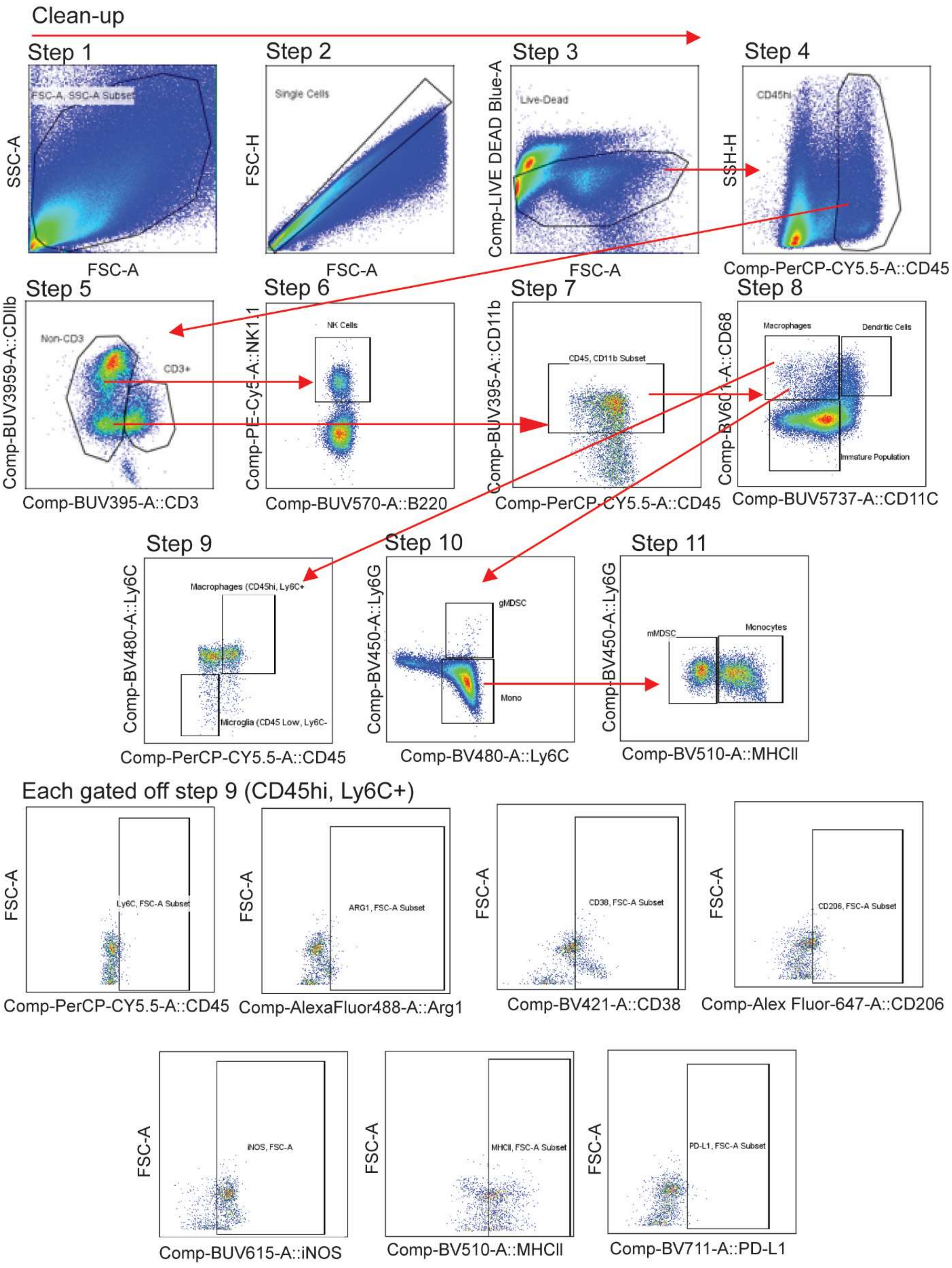


**Supplementary Figure 14. Tumor-infiltrating myeloid subsets flow cytometry clean up.** Gating strategy for the analysis of macrophage subsets in male and female mice treated with BMS986120 relative to vehicle.

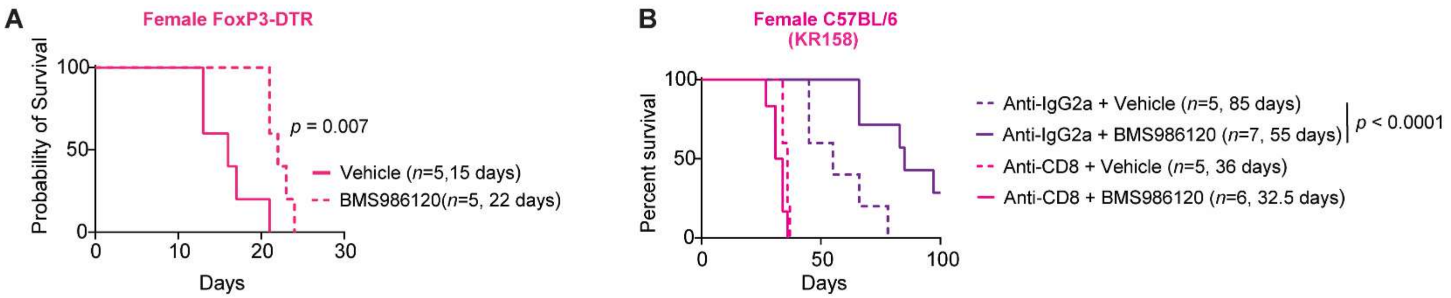


**Supplementary Figure 15. BMS986120 improves survival in C57BL/6 FOXP3-DTR mice. A.** Kaplan-Meier survival analysis was performed after intracranial implantation of the mouse GBM cell line SB28 in immunocompetent C57BL/6 FOXP3-DTR followed by pharmacologically inhibiting the thrombin PAR4 signaling axis by administering BMS986120 (2 mg/kg), a selective PAR4 inhibitor. **B**. Kaplan-Meier survival curve depicting survival of KR158 tumor-bearing females treated with anti-CD8-depleting antibody and BMS986120 administration. Statistical significance for survival analysis was determined by log-rank test.

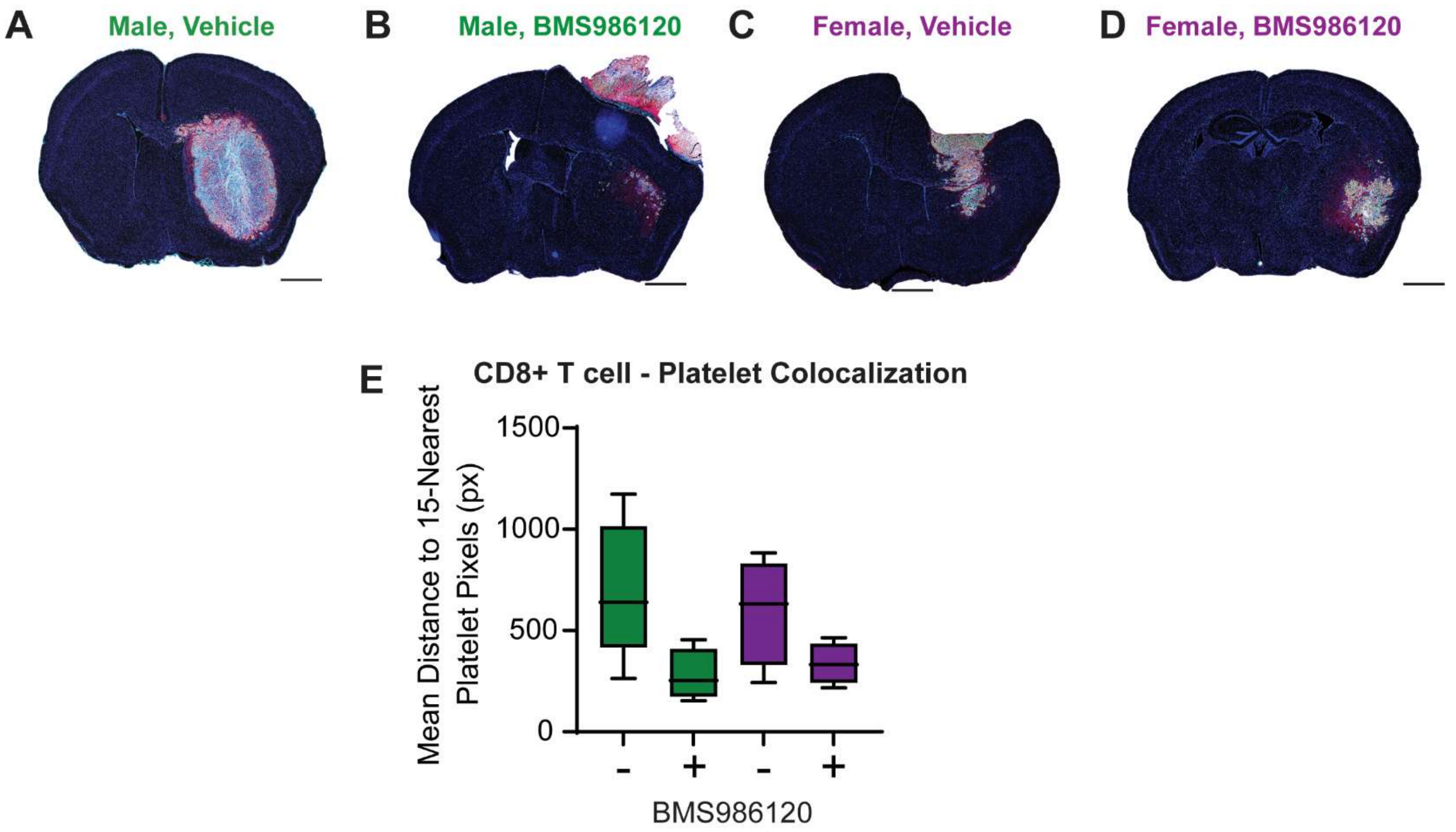


**Supplementary Figure 16. BMS986120 decreases intratumoral platelet expression and increase CD8+ T cells in females but not males**. **A-D**. Akoya multispectral immunofluorescence imaging analysis of in (**A**) male vehicle treated, (**B**) male BMS986120 treated, (**C**) female vehicle treated. **E**. Akoya multispectral immunofluorescence imaging analysis of the proximity of CD8+ T cells to platelet pixels.

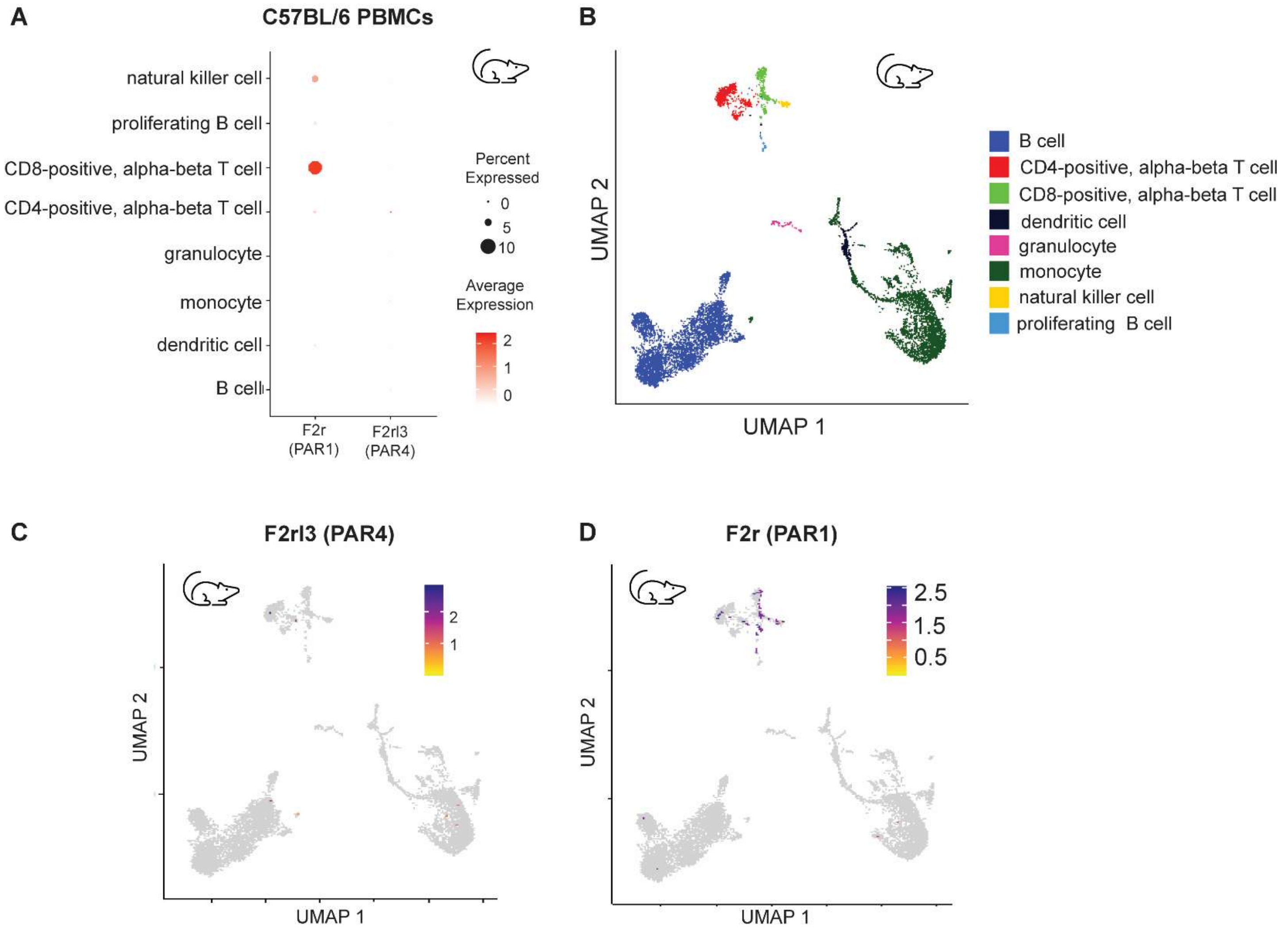


**Supplementary Figure 17. PAR1 but not PAR4 is expressed in mouse T cells. A.** FeaturePlots showing the average expression of PAR1 and PAR4 genes in PBMC samples from C57BL/6 mice. **B.** DotPlot showing the final cell clusters and markers to classify cells in mouse sc-RNAseq dataset. **C**, **D**. Dotplots showing the average expression of PAR4 (**C**) and PAR1 (**D**) in PBMC samples from C57BL/6 mice.

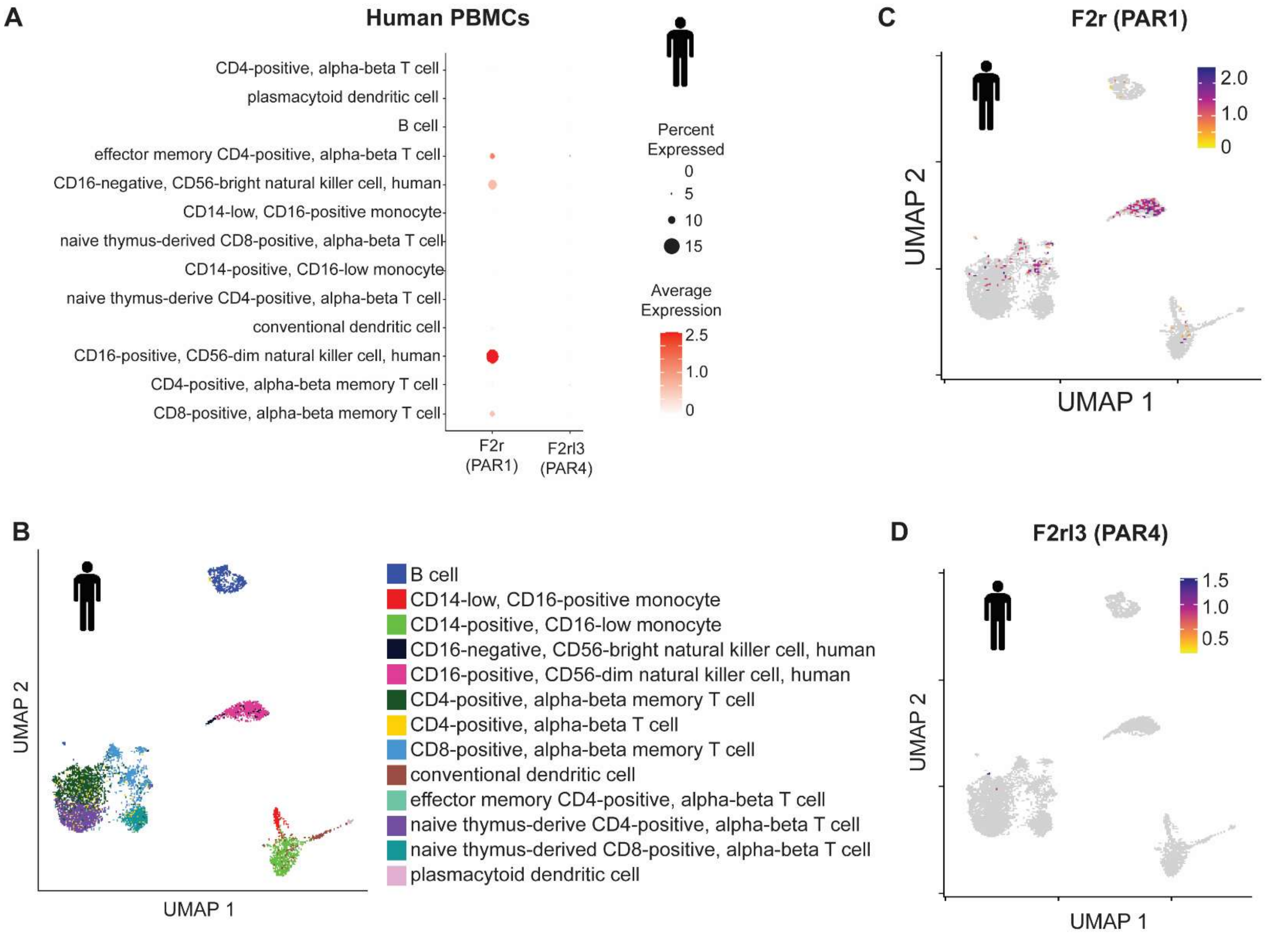


**Supplementary Figure 18. PAR1 but not PAR4 is expressed on human T Cells. A.** FeaturePlots showing the average expression of PAR1 and PAR4 genes in PBMC samples from healthy female subjects. **B.** DotPlot showing the final cell clusters and markers to classify cells in human sc-RNAseq dataset. **C**, **D**. Dotplots showing the average expression of PAR1 (**C**) and PAR4 (**D**) in PBMC samples from healthy female subjects.

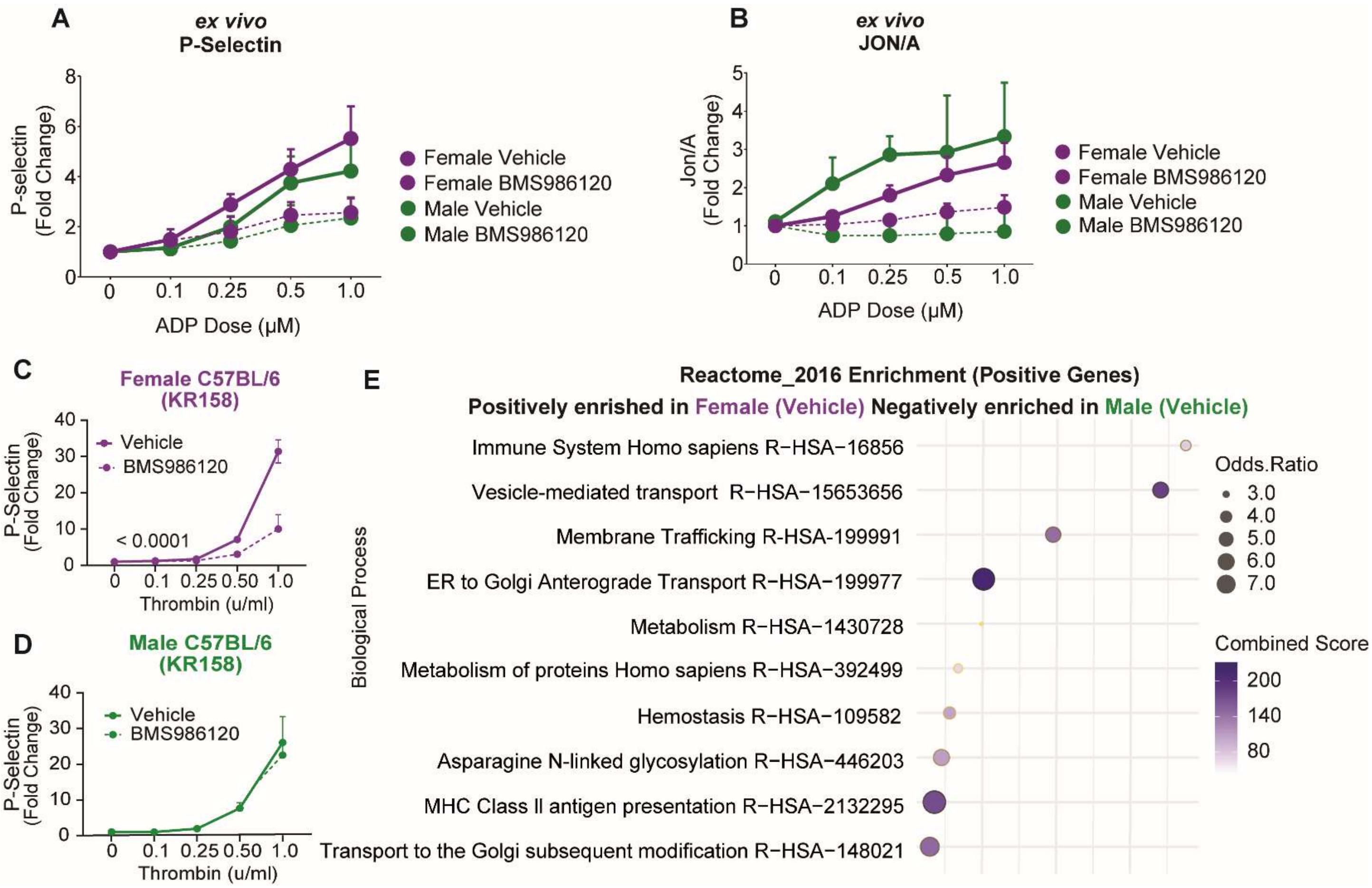


**Supplementary Figure 19. Sex biases between male and female platelets exist regardless of BMS986120 administration. A, B.** Washed platelets isolated from C57BL/6 mice treated with BMS986120 *ex vivo*. α-granule secretion was measured using an antibody specific for P-selectin (**A**), and activated GPIIb/IIIa was measured using the JON/A antibody (**B**). Data are represented as means ± SEM. Statistical significance was determined by two-way ANOVA. Washed platelets isolated from mice treated with BMS986120 following intracranial implantation of KR158 (**C, D**). Washed platelets were isolated, and α-granule secretion was measured using an antibody specific for P-selectin in female (**C**) and male (**D**) mice. Data are represented as means ± SEM. Statistical significance was determined by two-way ANOVA. **E**. Proteomic GO enrichment analysis of washed platelets isolated from mice treated with vehicle following intracranial implantation of SB28. Pathways are positively enriched in female platelets and negatively enriched in male platelets.

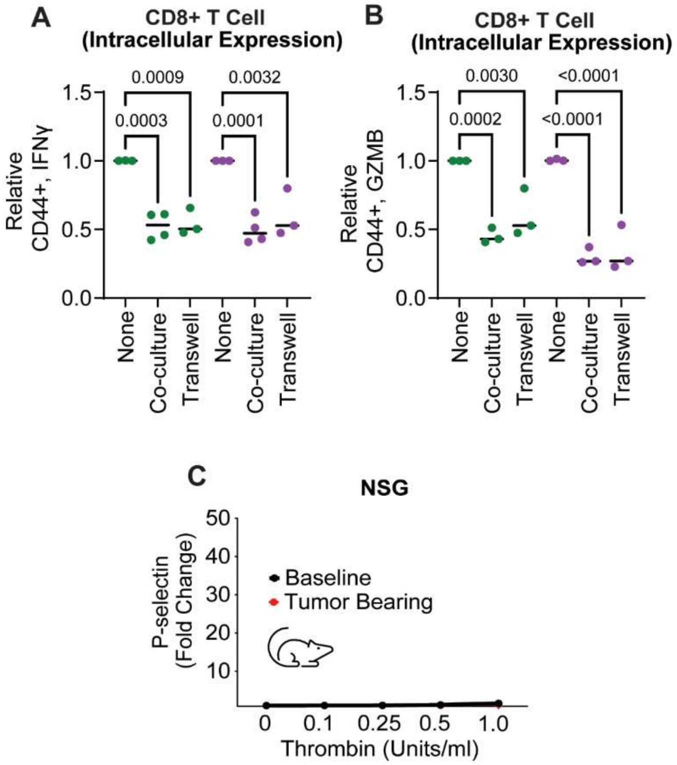


**Supplementary Figure 20. Tumor-bearing platelet reactivity in immune-incompetent mice. A, B**. Intracellular cytokine expression of CD8+ T cells isolated from C57BL/6 mouse spleen and co-cultured in the same well or through a transwell with sex-matched washed platelets isolated from C57BL/6 mice; fresh platelets were added for 3 consecutive days, and intracellular IFNγ (**A**) and GZMB (**B**) expression was measured by flow cytometry. One-way ANOVA analysis with Tukey's multiple comparison test was performed **C.** Washed platelets isolated from NSG mice before intracranial tumor implantation (baseline) and two weeks following intracranial implantation of SB28 cells. Washed platelets were isolated, and α-granule secretion was measured using an antibody specific for P-selectin via flow cytometry.

**Supplementary Table 1.** Platelet Gene Signature Genes identified by Kornblith et al., J Trauma Acute Care Surg., 2020.

| | | | |
|---|---|---|---|
| AKT3 | CSNK1G3 | IQGAP2 | PF4 |
| ANKRD12 | DNM3 | IQGAP2 | PLEK |
| ARHGAP18 | EIF4G3 | KIF2A | PTGS1 |
| ARHGAP21 | F13A1 | LIMS1 | RAP1B |
| ARHGEF12 | FIP1L1 | LTBP1 | RBPMS2 |
| CALD1 | FNBP1L | MAP4K5 | SH3BGRL2 |
| CCL5 | GRB14 | MBNL3 | SPARC |
| CDK2AP1 | HBA1 | NAP1L1 | TMSB4X |
| CHD9 | IQGAP2 | NLK | |

**Supplementary Table 2.** Platelet signature score refers to the Reactome pathway for ssGSEA

| | | | | | |
|---|---|---|---|---|---|
| A1BG | CD36 | FGA | LEFTY2 | PLG | SERPING1 |
| A2M | CD63 | FGB | LGALS3BP | PPBP | SOD1 |
| ABCC4 | CD9 | FGG | LHFPL2 | PPIA | SPARC |
| ACTN1 | CDC37L1 | FLNA | LY6G6F | PRKCA | SPP2 |
| ACTN2 | CFD | FN1 | MAGED2 | PRKCB | SRGN |
| ACTN4 | CFL1 | GAS6 | MANF | PRKCG | STX4 |
| AHSG | CHID1 | GTPBP2 | MMRN1 | PROS1 | STXBP2 |
| ALB | CLEC3B | HABP4 | NHLRC2 | PSAP | STXBP3 |
| ALDOA | CLU | HGF | OLA1 | QSOX1 | SYTL4 |
| ANXA5 | CTSW | HRG | ORM1 | RAB27B | TAGLN2 |
| APLP2 | CYB5R1 | HSPA5 | ORM2 | RARRES2 | TEX264 |
| APOA1 | CYRIB | IGF1 | PCDH7 | SCCPDH | TF |
| APOH | ECM1 | IGF2 | PCYOX1L | SCG3 | TGFB1 |
| APOOL | EGF | ISLR | PDGFA | SELENOP | |
| APP | ENDOD1 | ITGA2B | PDGFB | SELP | |
| BRPF3 | F13A1 | ITGB3 | PECAM1 | SERPINA1 | |
| CALM1 | F5 | ITIH3 | PF4 | SERPINA3 | |
| CALU | F8 | ITIH4 | PFN1 | SERPINA4 | |
| CAP1 | FAM3C | KNG1 | PHACTR2 | SERPINE1 | |
| CD109 | FERMT3 | LAMP2 | PLEK | SERPINF2 | |

**Supplementary Table 3.** Demographics for matched control subjects

| Study I.D | Sex | Age | Height (f'in”) | Weight (lbs) | BMI | Anti-platelet/Anti-coagulants |
|---|---|---|---|---|---|---|
| H1 | M | 46 | 5'10" | 200 | 26 | N/N |
| H2 | M | 45 | 6'0" | 200 | 25.8 | N/N |
| H3 | M | 33 | 6'0" | 175 | 22.5 | N/N |
| H4 | F | 44 | 5'10" | 160 | 20.9 | N/N |
| H5 | F | 37 | 5'9" | 150 | 22.2 | N/N |
| H6 | F | 37 | 5'6" | 110 | 17.8 | N/N |
| H7 | M | 54 | 6'0" | 220 | 29.8 | N/N |
| H8 | M | 55 | 5' 11" | 194 | 27.8 | N/N |
| H9 | M | 53 | 5'7" | 168 | 26.3 | N/N |
| H10 | F | 61 | 5'4" | 135 | 23.2 | N/N |
| H11 | F | 63 | 5'3 | 155 | 27.5 | N/N |
| H12 | F | 51 | 5'6 | 204 | 32.9 | N/N |
| H13 | F | 51 | 5'1.5" | 217 | 40.3 | N/N |
| H14 | F | 57 | 5'3'' | 190 | 33.7 | N/N |
| H15 | M | 55 | 6'0" | 220 | 29.8 | N/N |
| H16 | M | 54 | 5'7" | 168 | 26.3 | N/N |
| H17 | M | 56 | 5' 11" | 194 | 27.1 | N/N |
| H18 | F | 50 | 5'3" | 180 | 31.9 | Yes |

**Supplementary Table 4.** Demographics for GBM patients

| Study I.D | Sex | Age | Height (f'in") | Weight (lbs) | BMI | Anti-platelet/Anti-coagulants |
|---|---|---|---|---|---|---|
| G1 | F | 41 | 5'6" | 178 | 28.8 | N/N |
| G2 | F | 80 | 5'3" | 123 | 21.9 | N/N |
| G3 | F | 48 | 5'7" | 185 | 28.8 | N/N |
| G4 | F | 61 | 5'4" | 175 | 29.4 | N/N |
| G5 | F | 76 | 5'2" | 178 | 32.6 | N/N |
| G6 | M | 44 | 6'4" | 227 | 27.9 | N/N |
| G7 | F | 59 | 5'1" | 192 | 36.7 | N/N |
| G8 | F | 52 | 5'3" | 296 | 52.1 | N/N |
| G9 | M | 57 | 5'11" | 238 | 33.2 | N/N |
| G10 | F | 53 | 5'4" | 133 | 22.9 | N/N |
| G11 | F | 50 | 5'3" | 143 | 25.3 | N/N |
| G12 | M | 64 | 6'0" | 312 | 42.6 | N/N |
| G13 | M | 68 | 5'10" | 169 | 23.7 | N/N |
| G14 | F | 26 | 5'1" | 15 | 25.8 | N/N |
| G15 | M | 77 | 5'3" | 137 | 19.8 | Yes, Aspirin |
| G16 | M | 44 | 6'1 | 195 | 23.8 | N/N |
| G17 | M | 75 | 5'5" | 214 | 34.6 | Yes, Eliquis |
| G18 | M | 57 | 6" | 209 | 28.9 | N/N |

**Supplementary Table 5.** Flow antibodies for immune cell subsets

| Marker | Fluorophore | Vendor | Catalog number | Staining | Dilution |
|---|---|---|---|---|---|
| CD11b | buv395 | BD Biosciences | 563553 | Surface | 1:500 |
| CD69 | buv563 | BD Biosciences | 741234 | Surface | 1:500 |
| CD11c | buv737 | BD Biosciences | 612796 | Surface | 1:500 |
| CTLA4 | BV421 | BioLegend | 369606 | Intra | 1:250 |
| Ly6G | v450 | BioLegend | 127639 | Surface | 1:500 |
| PD1 | BV510 | BioLegend | 123118 | Surface | 1:250 |
| CD45R/B220 | BV570 | BioLegend | 103237 | Surface | 1:250 |
| Ki67 | BV605 | BioLegend | 652413 | Intra | 1:250 |
| TIM3 | BV711 | BioLegend | 300464 | Intra | 1:100 |
| CD3 | BV786 | Fisher Scientific | BDB564379 | Surface | 1:250 |
| CD45 | PerCP-Cy5.5 | Fisher Scientific | BDB561868 | Surface | 1:250 |
| Foxp3 | PE | Invitrogen | 12-5773-82 | Intra | 1:250 |
| LAG3 | PE-dazzle 594 | BioLegend | 125224 | Surface | 1:250 |
| NK1.1 | PE-Cy5 | BioLegend | 108716 | Surface | 1:250 |
| CD4 | PE-Cy7 | BioLegend | 100422 | Surface | 1:250 |
| CD8 | APC | BioLegend | 100712 | Surface | 1:250 |
| CD206 | Alexa Fluor 647 | BioLegend | 141712 | Intra | 1:250 |
| Ly6C | Alexa Fluor 700 | BioLegend | 128024 | Surface | 1:250 |
| CD68 | APC-Cy7 | BioLegend | 137024 | Intra | 1:250 |
| F4/80 | APC-Cy7 | BioLegend | 123128 | Surface | 1:250 |

**Supplementary Table 6.** Flow antibodies for cytokine profiling

| Marker | Fluorophore | Vendor | Catalog number | Staining | Dilution |
|---|---|---|---|---|---|
| CD45 | PerCP-Cy5.5 | BioLegend | 103132 | surface | 1:500 |
| CD3e | BV786 | BD Biosciences | 564379 | surface | 1:500 |
| CD8a | PE-Cy5 | BD Biosciences | 553034 | surface | 1:500 |
| CD4 | V450 | BD Biosciences | 560468 | surface | 1:500 |
| Foxp3 | Alexa Fluor 488 | eBioscience | 53-5773-82 | Intra | 1:250 |
| CD44 | BUV737 | BD Biosciences | 612799 | surface | 1:500 |
| IFN-γ | PE-dazzle 594 | BioLegend | 505846 | Intra | 1:250 |
| TNFα | BV605 | BioLegend | 506329 | Intra | 1:250 |
| Granzyme B | BV421 | BioLegend | 396414 | Intra | 1:250 |

**Supplementary Table 7.** Flow antibodies for myeloid immune cell subset profiling

| Marker | Fluorophore | Vendor | Catalog number | Staining | Dilution |
|---|---|---|---|---|---|
| CD11b | buv395 | BD Biosciences | 563553 | Surface | 1:500 |
| CD69 | buv563 | BD Biosciences | 741234 | Surface | 1:500 |
| iNOS | buv615 | Life Technologies | 366-5920-82 | Intra | 1:250 |
| CD11c | buv737 | BD Biosciences | 612796 | Surface | 1:500 |
| CD86 | buv805 | BD Biosciences | 741946 | Surface | 1:500 |
| CD38 | bv421 | BioLegend | 102732 | Surface | 1:500 |
| Ly6G | v450 | BioLegend | 127639 | Surface | 1:500 |
| Ly6C | bv480 | BioLegend | 128024 | Surface | 1:500 |
| MHCII | bv510 | BioLegend | 107636 | Surface | 1:500 |
| B220 | bv570 | BioLegend | 103237 | Surface | 1:500 |
| CD68 | bv605 | BioLegend | 137021 | Intra | 1:250 |
| PD-L1 | bv711 | BioLegend | 124319 | Surface | 1:500 |
| CD3 | bv786 | ThermoFisher | 417-0038-42 | Surface | 1:500 |
| Arg-1 | AF488 | SantaCruz Biotech | sc-271430 AF488 | Intra | 1:250 |
| CD8a | RB545 | Fisher Scientific | BDB569278 | Surface | 1:500 |
| CD45 | Percp-Cy5.5 | BioLegend | 103132 | Surface | 1:500 |
| *Plexin B2 | PE | BioLegend | 145903 | Surface | 1:500 |
| Ki-67 | PE-dazzle594 | BioLegend | 350534 | Intra | 1:250 |
| NK1.1 | PE-Cy5 | BD Biosciences | BDB561111 | Surface | 1:500 |
| CD4 | PE-Cy7 | BioLegend | 100422 | Surface | 1:500 |
| EGR2 | APC | eBioscience | 17-6691-80 | Surface | 1:500 |
| CD206 | AF647 | BioLegend | 141712 | Surface | 1:500 |
| F4/80 | APC-Cy7 | BioLegend | 123118 | Surface | 1:500 |

**Supplementary Table 8.** Immunofluorescence panel for Akoya multispectral immunofluorescence

| Marker | Clone ID | Vendor | Barcode | Reporter Dye | Source |
|---|---|---|---|---|---|
| c-Caspase 3 | 5A1E | Cell Signaling Technology | BX034 | Alexa Fluor™ 750 | Custom |
| c-KIT | YR145 | Abcam | BX331 | Alexa Fluor™ 750 | Custom |
| CCR7 | EPR23192-57 | Abcam | BX500 | Alexa Fluor™ 750 | Custom |
| CD11b | AKYP0149 | Akoya Biosciences | BX524 | Alexa Fluor™ 647 | Human IO |
| CD11c | AKYP0130 | Akoya Biosciences | BX024 | Alexa Fluor™ 647 | Mouse IO |
| CD163 | EPR19518 | Abcam | BX207 | Atto 550 | Custom |
| CD20 | AKYP0129 | Akoya Biosciences | BX064 | Alexa Fluor™ 750 | Mouse IO |
| CD206 | AKYP0137 | Akoya Biosciences | BX016 | Alexa Fluor™ 647 | Mouse IO |
| CD278 (ICOS) | AKYP0090 | Akoya Biosciences | BX065 | Alexa Fluor™ 647 | Human IO |
| CD304 | EPR3113 | Abcam | BX165 | Alexa Fluor™ 647 | Custom |
| CD31 | AKYP0131 | Akoya Biosciences | BX001 | Alexa Fluor™ 750 | Mouse IO |
| CD36 | AKYP0134 | Akoya Biosciences | BX002 | Atto 550 | Mouse IO |
| CD39 | AKYP0107 | Akoya Biosciences | BX099 | Atto 550 | Human IO |
| CD3e | AKYP0128 | Akoya Biosciences | BX080 | Alexa Fluor™ 647 | Mouse IO |
| CD4 | AKYP0146 | Akoya Biosciences | BX003 | Alexa Fluor™ 647 | Mouse IO |
| CD42b | polyclonal | | BX076 | Alexa Fluor™ 647 | Custom |
| CD44 | AKYP0132 | Akoya Biosciences | BX005 | Atto 550 | Mouse IO |
| CD45 | AKYP0143 | Akoya Biosciences | BX021 | Atto 550 | Mouse IO |
| CD45R/B220 | AKYP0014 | Akoya Biosciences | BX010 | Alexa Fluor™ 750 | Mouse IO |
| CD61/CD41 | Leo.A1 | | BX070 | Alexa Fluor™ 647 | Custom |
| CD62P | polyclonal | | BX518 | Alexa Fluor™ 647 | Custom |
| CD68 | AKYP0142 | Akoya Biosciences | BX015 | Alexa Fluor™ 647 | Mouse IO |
| CD8 | AKYP0145 | Akoya Biosciences | BX026 | Atto 550 | Mouse IO |
| Col1A1 | AKYP0141 | Akoya Biosciences | BX042 | Alexa Fluor™ 647 | Mouse IO |
| COL6A1 | EPR17072 | Abcam | BX116 | Alexa Fluor™ 750 | Custom |

| | | | | | |
|---|---|---|---|---|---|
| CXCR4 | UMB2 | Abcam | BX045 | Alexa Fluor™ 647 | Custom |
| F4/80 | AKYP0135 | Akoya Biosciences | BX006 | Alexa Fluor™ 647 | Mouse IO |
| FCRγ | AKYP0138 | Akoya Biosciences | BX037 | Alexa Fluor™ 750 | Mouse IO |
| FOXP3 | AKYP0133 | Akoya Biosciences | BX031 | Alexa Fluor™ 647 | Mouse IO |
| Histone H3 | AKYP0060 | Akoya Biosciences | BX030 | Alexa Fluor™ 647 | Human IO |
| Iba1 | AKYP0140 | Akoya Biosciences | BX014 | Atto 550 | Mouse IO |
| iNOS | AKYP0104 | Akoya Biosciences | BX023 | Atto 550 | Human Neuro |
| Ki67 | AKYP0052 | Akoya Biosciences | BX047 | Atto 550 | Mouse IO |
| Ly6g | AKYP0136 | Akoya Biosciences | BX017 | Alexa Fluor™ 647 | Mouse IO |
| MMP9 | RM1020 | Abcam | BX110 | Alexa Fluor™ 647 | Custom |
| P2RY12 | EPR26298-93 | Abcam | BX266 | Atto 550 | Custom |
| PCNA | AKYP0085 | Akoya Biosciences | BX036 | Alexa Fluor™ 750 | Human IO/Neuro |
| PDGFRB | Y92 | Abcam | BX049 | Alexa Fluor™ 750 | Custom |
| S100A10 | EPR3317 | Abcam | BX134 | Alexa Fluor™ 647 | Custom |
| S100A9 | AKYP0144 | Akoya Biosciences | BX025 | Alexa Fluor™ 750 | Mouse IO |
| SMA | AKYP0081 | Akoya Biosciences | BX013 | Alexa Fluor™ 750 | Human IO/Neuro |
| SOX2 | AKYP0106 | Akoya Biosciences | BX102 | Atto 550 | Human IO |
| TCF-1 | AKYP0099 | Akoya Biosciences | BX061 | Alexa Fluor™ 647 | Human IO |
| Ter119 | AKYP0013 | Akoya Biosciences | BX004 | Alexa Fluor™ 750 | Mouse IO |
| TOX | AKYP0098 | Akoya Biosciences | BX060 | Atto 550 | Human IO |
| VCAN | EPR12277 | Abcam | BX108 | Alexa Fluor™ 750 | Custom |
| Vimentin | AKYP0139 | Akoya Biosciences | BX022 | Alexa Fluor™ 750 | Mouse IO |
